# Targeting and combating drug resistance in triple-negative breast cancer using nano polymer: Efficacy of A6 peptide-PLGA-PEG nanoparticle loaded with doxorubicin and anti-miR-21 in *in vitro* and *vivo model*

**DOI:** 10.1101/2025.01.04.631287

**Authors:** Zaidah Ibrahim, Pei Pei Chong, Hasan Al-Moustafa, Maizaton Atmadini Abdullah, Zamri Chik

## Abstract

Triple-negative breast cancer (TNBC) poses a significant challenge in treatment due to the emergence of drug resistance to standard chemotherapeutics, along with associated risks to non-targeted tissues. This study is dedicated to developing a nanoparticle drug delivery system utilizing PLGA-PEG nanoparticles (NP) with the integration of A6 peptide (NPA6) as a specific targeting ligand for TNBC. Encapsulating Doxorubicin (DOX) and antisense-miR-21 (AM21), these nanoparticles are designed to counter drug resistance in vitro and in vivo. Characterized by an average size of 103.5 (±8.1) nm and remarkable DOX encapsulation efficiency, these nanoparticles have been successfully engineered. Results demonstrate that NPA6.DOX.AM21 effectively mitigates drug resistance, displaying a significantly decreased doxorubicin IC_50_ as opposed to free DOX (2.5 µM vs 25.7µMrespectively, p<0.01). In mice models, the A6 peptide-incorporated formulation (NPA6.DOX, NPA6.DOX.AM21) exhibits substantial tumour size reduction and increased doxorubicin concentration within tumours, all while minimizing non-specific doxorubicin exposure compared to non-A6 formulations (NPDOX, FREE DOX) treatments. This innovative PLGA-PEG-A6 peptide complex loaded with DOX and AM21 showcases superior efficacy in surmounting cell resistance in breast cancer cell line, targeting tumour progression, and decreasing drug distribution in non-targeted organs in mice. These results hold promise for enhanced combat against drug resistance and precise chemotherapy, hence potentially improving patient outcomes and survival rates.

**Graphical abstact:** 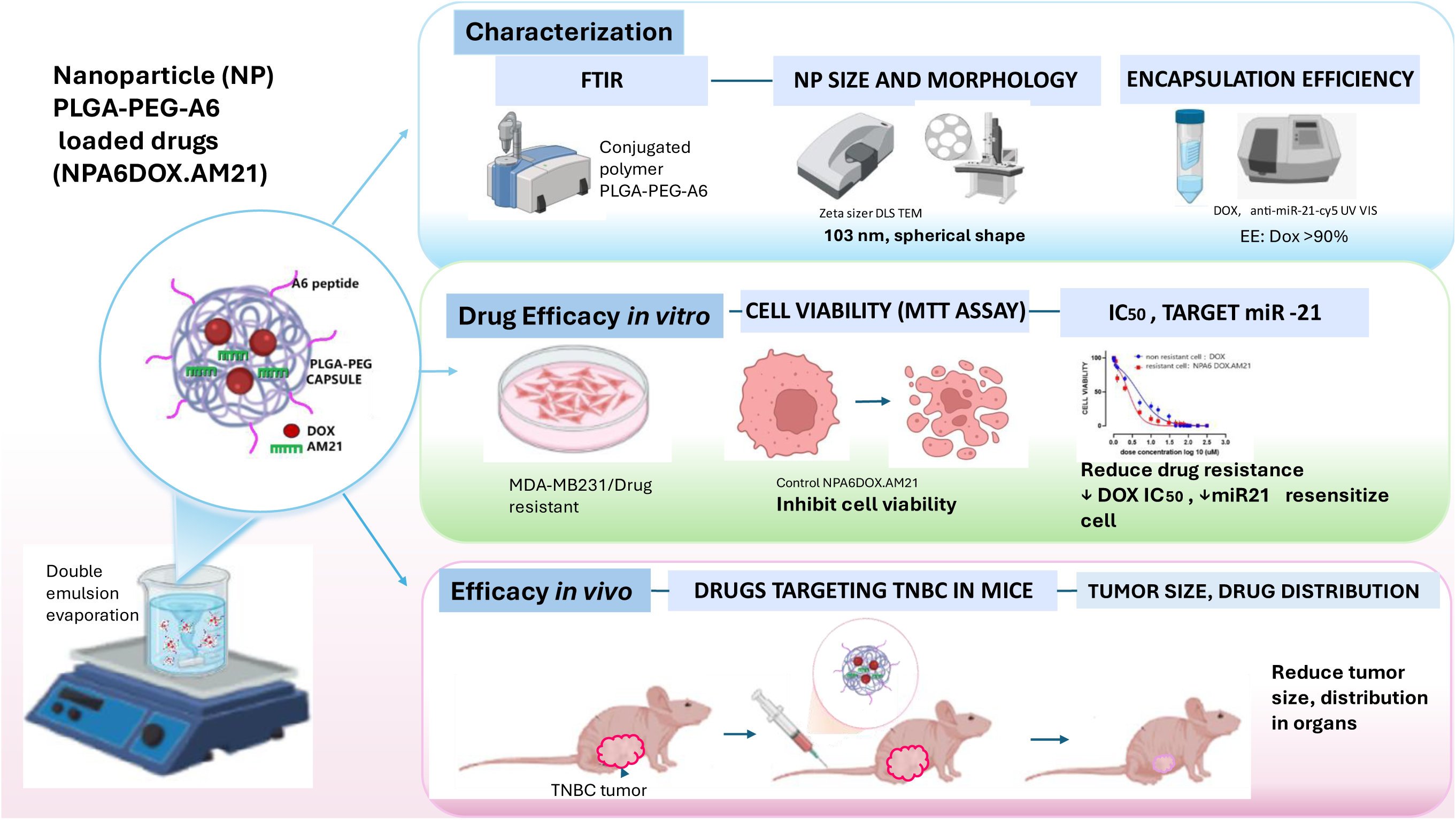

Figure Graphical abstract of the study. Fron left to right, development of nanoparticles formulation a followed by characterization and efficacy of the nanoparticle formulation in *in vitro* and *in vivo* to reduce drug resistance and target triple negative breast cancer (TNBC). PLGA: polylactic-co-glycolic acid, PEG: polyethylene glycol, A6: A6 peptide, DOX: Doxorubicin, AM21: antisense-microRNA -21

**Highlights:**

- The nanoparticle of A6 conjugated PLGA-PEG co-loaded Doxorubicin with anti-miRNA21 (NPA6.DOX.AM21) was successfully fabricated with the size of 99.8 - 103.5 nm diameter and PDI 0.2 with high drugs encapsulation efficiency.
- NPA6.DOX.AM21 increased cell death, reduced drug resistance in MDA MB-231 resistant cells, and reduced IC_50_ of doxorubicin dose.
- NPA6.DOX.AM21 targeted and reduced TNBC tumour size, with high doxorubicin distribution in tumor and less accumulation in off-target organs, in comparison to the control in mice model.

## 1 Background

Triple-negative breast cancer (TNBC) represents a highly aggressive subtype of breast cancer distinguished by the absence of oestrogen receptor (ER), progesterone receptor (PR), and human epidermal growth factor receptor 2 (HER2) expression. Owing to the absence of these receptors, TNBC exhibits resistance to hormonal therapies and targeted treatments, posing a significant clinical challenge due to the lack of specific therapeutic options available [1]. In TNBC, the preferred chemotherapy agent is doxorubicin (DOX), renowned for its potent DNA chelating properties that impede cell cycle progression, induce apoptosis, and exhibit high cytotoxicity against cancer cells. Despite its efficacy, the current formulation of doxorubicin lacks ideal attributes as TNBC tumours frequently develop resistance to the drug post-treatment, resulting in cancer relapse [2].

Doxorubicin is employed in the treatment of various cancers. However, its clinical utility is limited by its adverse effects on essential organs, including the heart and bone marrow. Cardiotoxicity associated with doxorubicin usage can lead to long-term cardiovascular complications in cancer survivors [3]. Therefore, there exists a crucial necessity to enhance the targetability and efficacy of doxorubicin while minimizing its detrimental effects.

Studies have shown that tumour proliferation and the development of drug resistance are influenced in part by microRNAs (miRs), which can modulate tumor-suppressing proteins [4]. Among these miRs, miR-21 has garnered significant attention due to its overexpression in multiple cancer types, including TNBC [5]. Notably, miR-21 has been implicated in conferring resistance to doxorubicin in TNBC [6] and gastric cancer [7]. Its overexpression has been correlated with heightened cell growth, metastasis, and drug resistance in diverse cancer types. Inhibiting miR-21 using antisense miR-21 molecules has shown promise in reducing cancer cell growth and metastatic potential [6]. Functionally, miR-21 exerts its effects by targeting tumour suppressor proteins, facilitating cancer cell invasion and metastasis [8]. Targets of miR-21 such as PTEN and PDCD4 have been associated with modulating breast cancer cell growth and inducing resistance to doxorubicin [9]. Additionally, miR-21 has been linked to drug [8] resistance in various cancers such as gastric, colon, nasopharyngeal, and prostate cancer by targeting PTEN and other key molecules, underscoring its significant role in cancer progression and resistance [10][11][12][13]. Combining miR-21 inhibition with doxorubicin has exhibited enhanced efficacy against MCF7 cells, illustrating increased drug susceptibility [14]. However, investigations into the combination of doxorubicin with anti-miR-21 to combat drug resistance in TNBC models, specifically the MDA-MB-231 cell line, remains unexplored.

Moreover, doxorubicin chemotherapy in its conventional formulation may cause severe toxic effects on non-target organs, leading to poor survival rates and prognoses in TNBC patients. The drug not only eradicates cancer cells but also accumulates in the heart muscle, resulting in cardiotoxicity [15]. To address this limitation, there is a growing focus on utilizing drug-loaded targeted nanocarriers to improve drug delivery. Owing to their nanoscale size, nanocarriers can traverse compromised cancer site endothelial barriers and augment the delivery of anticancer agents to tumours [16]. Furthermore, these nanocarriers can be functionalized with specific molecules to selectively target cancer cells [17].

Among the innovative drug carriers, nano-polymer micelles composed of PLGA-b-PEG have emerged as US FDA-approved entities with high bioavailability and biocompatibility. The presence of PEG molecules acts as a protective shield against enzymatic degradation upon entering the circulatory system [18]. PLGA-PEG can be modified with diverse functional groups to enable conjugation with specific targeting molecules, facilitating targeted drug delivery strategies [19]

Transmembrane receptor CD44 overexpression is commonly observed in many malignant cancers, including triple-negative breast cancer (TNBC), and is associated with increased cell signalling and tumour metastasis. In TNBC MDA-MB-231 cell line, CD44 is prominently expressed on the cell surface [20], providing a potential target site for ligand-receptor interaction. Recent advancements have demonstrated the potential of using peptides as targeting molecules to bind specifically to cells expressing particular surface receptors, enhancing drug delivery. Studies have indicated that the A6 peptide, which shares a partial homology with the CD44 binding site, can effectively target cancer cells with minimal side effects [20][21][22][23].

Considering the specific binding between the A6 peptide and CD44, incorporating the A6 peptide onto PLGA-PEG nanocarriers as a cancer-targeting element holds significant promise. However, the potential of the PLGA-PEG polymer to encapsulate doxorubicin along with other anti-cancer agents such as anti-miR-21 and conjugating it with the A6 peptide for targeted delivery in TNBC is still largely unexplored. It is essential to verify in experimental studies the efficacy of this new formulation in navigating the host circulatory system to reach the tumor site effectively while minimizing accumulation in non-target organs like the heart.

To date, there is a lack of research utilizing the PLGA-PEG formulation conjugated with the A6 peptide and loaded with doxorubicin in combination with anti-miR-21 to address drug resistance and non-specific targeting in the resistant MDA-MB-231 cell model both *in vitro* and *in vivo*. It is crucial to develop an innovative formulation to tackle the challenges encountered in TNBC chemotherapy.

In this study, we successfully synthesized PLGA-PEG nanoparticles conjugated with A6, loaded with doxorubicin (DOX) and anti-miR-21 (AM21) (NPA6DOX.AM21) and assessed its efficacy against resistant cells, as well as examined the A6 peptide targeting effect in the TNBC model both *in vitro* and *in vivo*. The study hypothesized that the synthesized NPA6.DOX.AM21 would exhibit favourable nanoparticle characteristics, overcome doxorubicin resistance in MDA-MB231 cells, enhance drug delivery to the tumour, and reduce tumour size in TNBC xenografted mice.

## 2. Methodology

### 2.1 Development of nanoparticle drug

#### Material

Poly (D, L-lactide-co-glycolide) (in short PLGA) (LA:GA ratio 50/50) with carboxylic acid terminated (PLGA-COOH, inherent viscosity 0.16-0.24 dL/g, 8000mw) (from LACTEL. EVONIK brand). EDC (1-ethyl-3-(3-(dimethylamino) propyl) carbodiimide, N-hydroxy succinimide (NHS), 4 dimethyl aminopyridine (DMAP), and diisopropylethylamine (DIPEA), were purchased from Sigma Aldrich. Polyethylene glycol NH_2_-PEG-COOH (PEG2000), (from BIOCHEM PEG brand), triethylamine (TEA), Doxorubicin (DOX.HCL) mw 580 g/mol. A6 peptide with sequence of Ac-KPSSPPEE-NH2, nucleic acid materials: miR-21-*S, Anti-miR-21-*S, Cy5-Anti-miR-21-*S, Anti-sense-MiR-21 (mw 6810 mg/mmol, 4.6 nmol/0D 260, 31.1 ug/OD260) were obtained from IDT Technology, Singapore. Filter membrane ultracentrifugation tubes 10,000 mw pore (Amicron brand). Solvents: DCM, methanol, acetonitrile (ACN), diethyl ether, (Sigma Aldrich) sterile distilled water, nuclease enzyme-free deionized water. (Elga water system)

#### 2.1.1 Synthesis of nanoparticles

The synthesis of nanoparticles involved the preparation of PLGA-PEG and PLGA-PEG-A6, followed by the encapsulation of the anticancer agents. The coupling of PLGA to PEG (PLGA-PEG) and PLGA-PEG to A6 peptide (PLGA-PEG-A6) was achieved using the EDC/NHS reaction to establish an amid bond linkage between the two materials. The confirmation of the PLGA-PEG and PLGA-PEG-A6 coupling synthesis was conducted through FTIR analysis.

##### Drug encapsulation

Eight nanoparticle (np) formulations were meticulously prepared, consisting of 6 treatment np formulations loaded with anticancer agents, and 2 nanoparticle (np) formulations without anticancer agents serving as controls (blank np).

The formulations of NP and NPA6 (NP: PLGA-PEG), (NPA6: PLGA-PEG-A6) were loaded with either single doxorubicin (DOX) or single anti-sense miR-21 (AM21) or combination of DOX and AM21 (DOX.AM21) **(Table 1).**

**Table 1.** Nanoparticle formulation.

|  | Formulation with NP<br>(PLGA-PEG) | Formulation with NPA6<br>(PLGA-PEG-A6) |
| --- | --- | --- |
| *(blank)* | NP | NPA6 |
| *Single loaded* | NPDOX | NPA6 DOX |
| *Single loaded* | NPAM21 | NPA6AM21 |
| *Co-loaded* | NPDOX.AM21 | NPA6DOX.AM21 |

DOX: doxorubicin , AM21: anti-miR21

The encapsulation of doxorubicin and anti-miR21 was carried out using the double emulsion solvent evaporation method with specific modifications adapted from Devulapaly et al., 2015 and Mosafer et al., 2018 [24] [25] as depicted in Fig. 2 and supplementary information <u>S.Info</u>. Subsequently, the nanoparticles were filtered and dried.

**Fig. 1.**
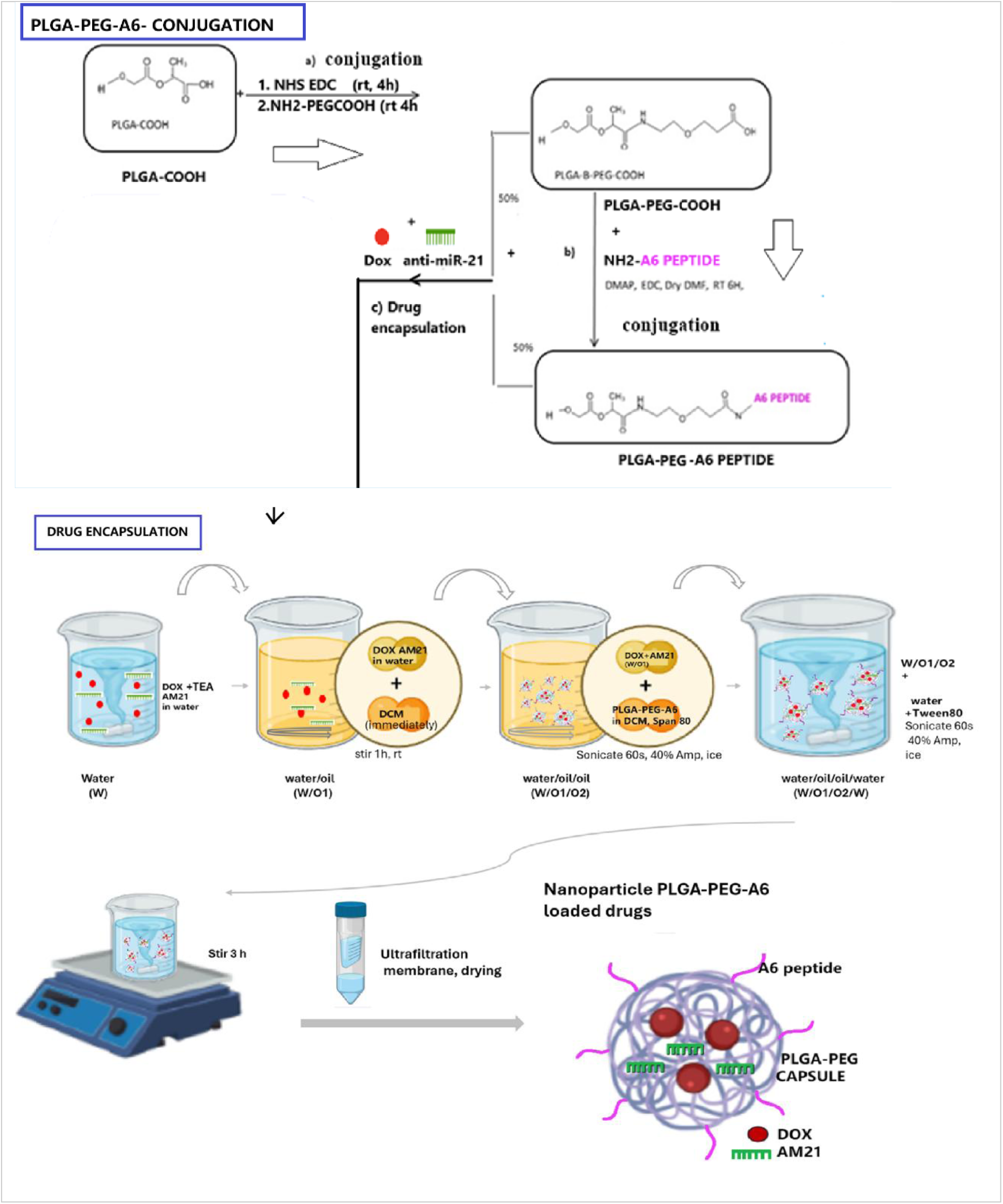
Schematic illustration of nanoparticle synthesis. From top to bottom. Conjugation of a. PLGA and PEG into PLGA-PEG and b. PLGA-PEG and A6 into PLGA-PEG-A6. c. drug encapsulation using double emulsion and solvent evaporation method co-encapsulating DOX and AM21 in PLGA-b-PEG and PLGA-PEG-A6 producing polymer nanoparticle loaded drugs. DOX: doxorubicin, AM21: anti-sense micro-RNA21

**Fig. 2.**
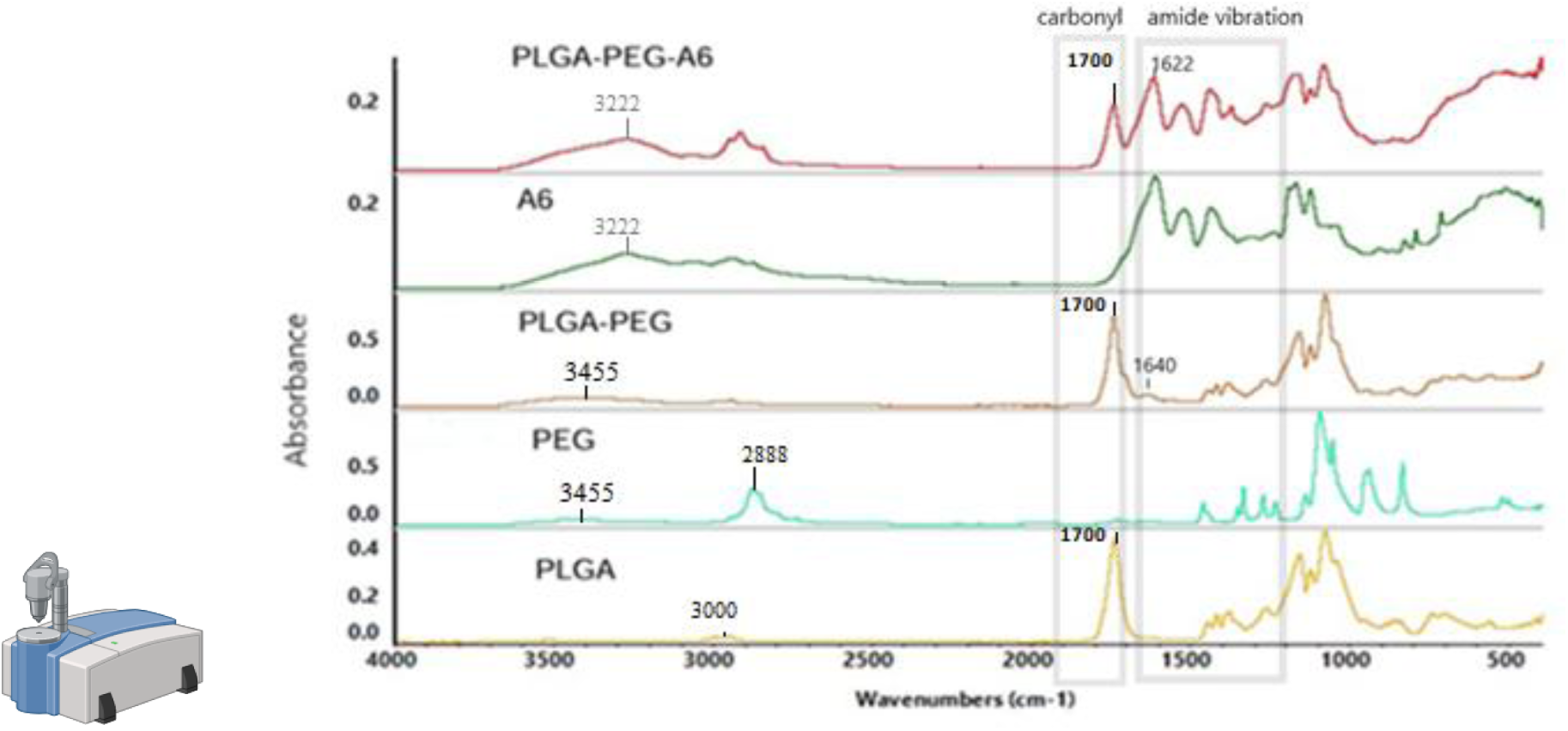
FTIR analysis of PLGA-PEG and PLGA-PEG-A6 conjugation. The box indicates carbonyl and amide vibration band shift when new amide bond was formed. Absorbance intensity of band shifts between 1500 to 1700 cm^-1^ were noted. PLGA-COOH carbonyl compound peak detected at 1700 cm^-1^, while NH2 from PEG-NH2 detected at 3455cm^-1^ individually. After conjugation, the carbonyl stayed at 1700cm^-1^ in PLGA-PEG while new amide peak appeared at 1640 cm^-1^ compared to PLGA and PEG respectively. A6 showed N-H stretch between 3170 to 3500 cm^-1^ and amide vibration between 1500cm^-1^ to 1700cm^-1^. Following PLGA-PEG conjugation to A6 peptide, similar spectrum to PLGA-PEG and A6 were detected with new peak appeared at 1622 cm^-1^ as well as at 3222 cm^-1^ confirming the successful conjugation of the PLGA-PEG-A6 polymer chain.

Characterization of the np formulation was performed by measuring size, zeta potential, and polydispersity index (PDI) through dynamic light scattering (DLS) and Zetasizer equipment (Malvern Panalytical). The morphology of the polymer was evaluated by staining the nanoparticles with 1% (v/v) phosphotungstic acid and observing them using TEM microscopy. The entrapment of Dox and AM21 in the np was quantified by absorbance of Dox and AM21 using UV-VIS spectrophotometer (Perkin Elmer).

### 2.2 Evaluation of drug resistance in cells

The development of drug-resistant cells involved transfecting cells with the plasmid (pCI-neo Fluc EGFP was a gift from Franz-Ulrich Hartl (Addgene plasmid # 90170; http://n2t.net/addgene:90170; RRID:Addgene_90170) [26] containing ABCB1 resistance gene (cloned at GenScript, USA, Clone ID:OHu23307C) tagged with the reporter eGFP gene, resulting in green fluorescence cells when expressed. The cell transfection was performed using a transfection kit (Thermo Fisher Scientific). These resistance-gene transfected cells were repeatedly exposed to increasing concentrations of 1-8 µM Dox. Cells that survived at the highest concentration (8 µM) were isolated and cultured to maintain the drug-resistant phenotype.

#### 2.2.1 Evaluation on Cell viability and cytotoxicity IC50 on the cancer cells

To compare the effects of the np formulations on cell viability and IC_50_ values, both parental (non-transfected/non-resistant) cells and drug-resistant cells were subjected to free doxorubicin treatment. Cells were seeded in a 96-well plate with different concentrations of doxorubicin. Cell viability was assessed through the MTT assay, with mixing solutions prepared using MTT powder and PBS. MTT assay ([(3-(4,5-dimethylthiazol-2-yl)-2,5-diphenyl-tetrazolium bromide) was carried out at 24 h after exposure treatments to see the cell viability. Mixing solutions were prepared using 2 mg of MTT (Sigma-Aldrich) powder in 1 ml PBS. The culture medium was changed with 150 μl fresh media plus 50 μl MTT reagent (2 mg/ml in PBS); the cell-free wells were considered as blank controls. Cells were then incubated at 37°C with CO_2_ at 5% and a humidified atmosphere for 4 h. Then, the MTT solution was removed and 200 μl of DMSO was added to each well. The OD of the wells was measured at 550 nm using a spectrophotometric microplate reader to determine cell viability.

Following the initial comparison, the resistant cells were further tested using nanoparticle formulations and free doxorubicin at doxorubicin concentrations ranging from 0-100 µM. The impact of co-loading AM21 (25 pmol) with doxorubicin in the np formulation on cell viability and the reduction of the doxorubicin IC_50_ derived from dose response curve, were evaluated to determine the effectiveness of this combination in inhibiting cell growth

#### 2.2.2 Evaluation of the formulation on mi downstream tar et assessing miR-21, mRNA PTEN and PDCD4 level

The miR-21 level, and the mRNA of PTEN and PDCD4 level were measured in both MDA-MB-231 non-resistance cells (wild type/ non transfected with ABCB1 gene) and in transfected ABCB1 gene/ resistance cells (MDA-MB-231/R). Comparison between these two cell types would further confirm the changes level in resistant cells. This was to assess the basal level before the nanoparticle treatment given to the cells

Following the development of resistant cells, the MDA-MB-231/R resistant cells were treated with the prepared formulations containing 25 pmols of AM21 loaded np as well as the rest of the np formulations for 48 hours to see if the antisense miR-21 affected the expression level of its cellular miR-21, and its target mRNA PTEN and PDCD4 by RT-PCR.

##### RT-PCR

Total RNA from the treated and control cell samples were extracted using RNeasy kit (QIAGEN, Germany) according to the manufacturer’s instructions. The extracted RNA was then evaluated for expression level of mature miR-21, mRNA PTEN and PDCD4 as determined by the RT-PCR miRNA-assay (Applied Biosystems, Foster City, CA, USA). All RT-PCRs were performed in triplicates. The RT primer, PCR primers, for miR-21, PTEN and PDCD4 were purchased from local providers. The sequence of the primers was shown in supporting information. The real-time PCR results were analyzed and expressed as relative miRNA and mRNA expression of CT (threshold cycle) value, which were then converted to fold changes. The house keeping gene for miR-21 was normalized to reference gene U6 RNA and fold expression were calculated. While housekeeping gene β-actin were used for normalizing the gene expression of target mRNA PTEN and PDCD4.

### 2.3 Evaluation of nanoparticle formulation on TNBC tumour xenograft in mice model

Animal ethics research approval was obtained from the Institutional Animal Care and Use Committee (IACUC), Faculty of Medicine, Universiti Malaya (UM) (2023-240105/PHARM/R/HA). There were 5 groups of mice, each group consisting of 6 mice: a control group/blank NP, free doxorubicin, NP.DOX, NPA6.DOX, NPA6DOX.AM21.

#### 2.3.1 Establishment of tumor cell xenograft model

Female athymic Balb/c nu/nu mice, 6 weeks old (obtained from Charles River Laboratory, USA), were used as the tumour model for evaluating nanoparticle drug efficacy. The mice were housed in cages at 23 ± 2 °C with a 12-hour light/dark cycle and 55 ± 10% humidity, provided with standard mouse pellets and water ad libitum.

MDA-MB-231 cell line cultures were harvested, centrifuged at 1000 rpm for 3 min and then resuspended in 1:1 PBS and Matrigel mixtures at a density of 5×10^6^ cells/mL (adapted from [27]).

Each cell resuspensions containing 100 µL was subcutaneously injected to establish tumour on the lower flanks of the mice near the mammary pad. The mice were monitored for their tumour growth size measured using calliper using the formula, tumour volume= (length×width×width)/2 from [28]. Tumour growth was monitored using callipers until reaching a size of 100mm³ to commence treatment.

#### 2.3.2 Evaluating antitumor properties of nanoparticle formulation in TNBC tumor xenograft

At 100mm^3^ of tumor volume size, the mice were regrouped into 6 mice per group. The treatment starts at Day 0 as first injection and was given through tail vein injection. The dose was 100 ul PBS containing Dox formulation equivalent of 2.5 mg/kg for each 1^st^, 2^nd^ and 3^rd^ injection and 5 mg/kg for each 4^th^ and 5^th^ tail vein injection (Day 0, 5,10,15,20) (adapted from [29]). As for the control group, the mice received 200 uL of saline with empty NP. The tumor volume was continuously monitored until Day 24.

### 2.4 Biodistribution of nanoparticle formulation in tumor and off-target organs

To assess drug accumulation specificity in the tumor between A6-nanoparticle and non-A6 formulations, a dose equivalent to 10 mg/kg of dox-loaded formulation was injected via the tail vein on Day 23. After 24 hours, the mice were euthanized, and organs and tumors were harvested for Dox fluorescence imaging using an FX Pro Bioimaging instrument with 530/600 nm filters. Dox fluorescence intensity in tumors and organs was compared among treatment groups using ImageJ software. Higher Dox fluorescence intensity indicated greater accumulation of Dox in the tissues.

#### Tumor and organs histology (H&E staining)

Following bioimaging, tumors and organs were fixed in formalin, and fresh frozen FFPE blocks were prepared. Tissue sections of 5μm were stained with H&E to observe histomorphological changes.

### 2.5 Data analysis and statistics

Nanoparticle development and characterization were descriptively analysed. Efficacy testing of drug formulations in in vitro and in vivo models was subject to multi-comparison assessment between treatment groups using ANOVA followed by post-test analysis with Bonferroni correction. Further paired t-tests were conducted for two-group comparisons (within resistant cells) and unpaired independent t-test (between resistant and non-resistant cell) were conducted accordingly. Statistical significance was set at p<0.05.

## 3 Results and Discussion

### 3.1 Nanoparticle Characterization

The success of nanoparticle preparation evaluated by the PLGA-PEG and PLGA-PEG-A6 conjugation followed by the characterization of the nanoparticle loaded drug. The conjugation of PLGA-PEG and PLGA-PEG-A6 peptide was characterized by FTIR analysis, which confirmed the formation of amide bonds between PLGA and PEG, as well as between PLGA-PEG and the A6 peptide (Fig. 2). The synthesis reaction utilized carboxylic acid (-COOH) to link the carbonyl group of one component to the nitrogen atom of the next, forming the amide bond. As shown in Figure 2, PLGA-COOH displayed a characteristic carbonyl peak at 1700 cm⁻¹, along with a small doublet around 3000 cm⁻¹ (attributed to CH, CH2, and CH3). In contrast, PEG exhibited a prominent band at 2888 cm⁻¹ (CH, CH2, and CH3) and an N-H stretch at 3455 cm⁻¹ for PEG-NH2. Following the conjugation of PLGA to PEG, the spectra for PLGA and PEG showed similarities, but distinct changes were observed in the region from 1700 cm⁻¹ to 1500 cm⁻¹, with a new peak appearing at 1640 cm⁻¹, indicating amide formation, and the N-H stretch at 3455 cm⁻¹ was also noted. For clarity, the dotted box in Fig. 2 highlights the carbonyl and amide vibrations. The spectrum of the A6 peptide showed a notable N-H stretch between 3170 to 3500 cm⁻¹ and amide vibrations around 1500 cm⁻¹ to 1700 cm⁻¹. After the conjugation of PLGA-PEG to the A6 peptide, a similar spectrum was observed for both PLGA-PEG and A6, with the emergence of a new peak at 1622 cm⁻¹ reflecting amide vibration, as well as at 3222 cm⁻¹, indicative of the N-H stretch, signifying that amide bonds were formed and there was a loss of the -OH group from PLGA-PEG-COOH. The reference bands for identifying amide bonds through infrared absorption have been adapted from previous studies [30].

Following an encapsulation process, the PLGA-PEG (NP) and PLGA-PEG-A6 (NPA6) nanoparticles achieved a size close to 100 nm in diameter, with average sizes ranging from 99.8 to 103.5 nm, making them suitable for drug delivery. This size falls within the desired range of 100-200 nm for nanoparticles. Previous research has shown that nanoparticles below 50 nm had tendencies to accumulate in liver tissues, while those above 400 nm had difficulties entering tumor areas through the capillary membrane leakage, thus limiting the potential of the enhanced permeability and retention (EPR) effect. Ideally, nanoparticles for nano biomaterial applications related to the EPR effect should fall between 100 and 400 nm [31] which this current study may take advantage of.

The size of the nanoparticles developed in this study was confirmed to be around 100 nm in diameter under TEM microscopy (Fig. 3). Report shows that nanoparticles with sizes below 200 nm are typically effective in targeting and facilitating drug delivery to cancer cells. Particularly, size plays a crucial role in cell uptake, tissue penetration, and circulation time, with smaller nanoparticles having better chances of exhibiting these desired characteristics. Smaller particles can evade clearance in systemic circulation, thereby enhancing drug delivery efficiency [33]. In this study, the polydispersity index (PDI) (Table 2) of the nanoparticles was 0.2, indicating towards a monodisperse or a good size distribution inferring good effect of the nanoparticles delivery towards the cells [33] [34][35].

**Fig. 3.**
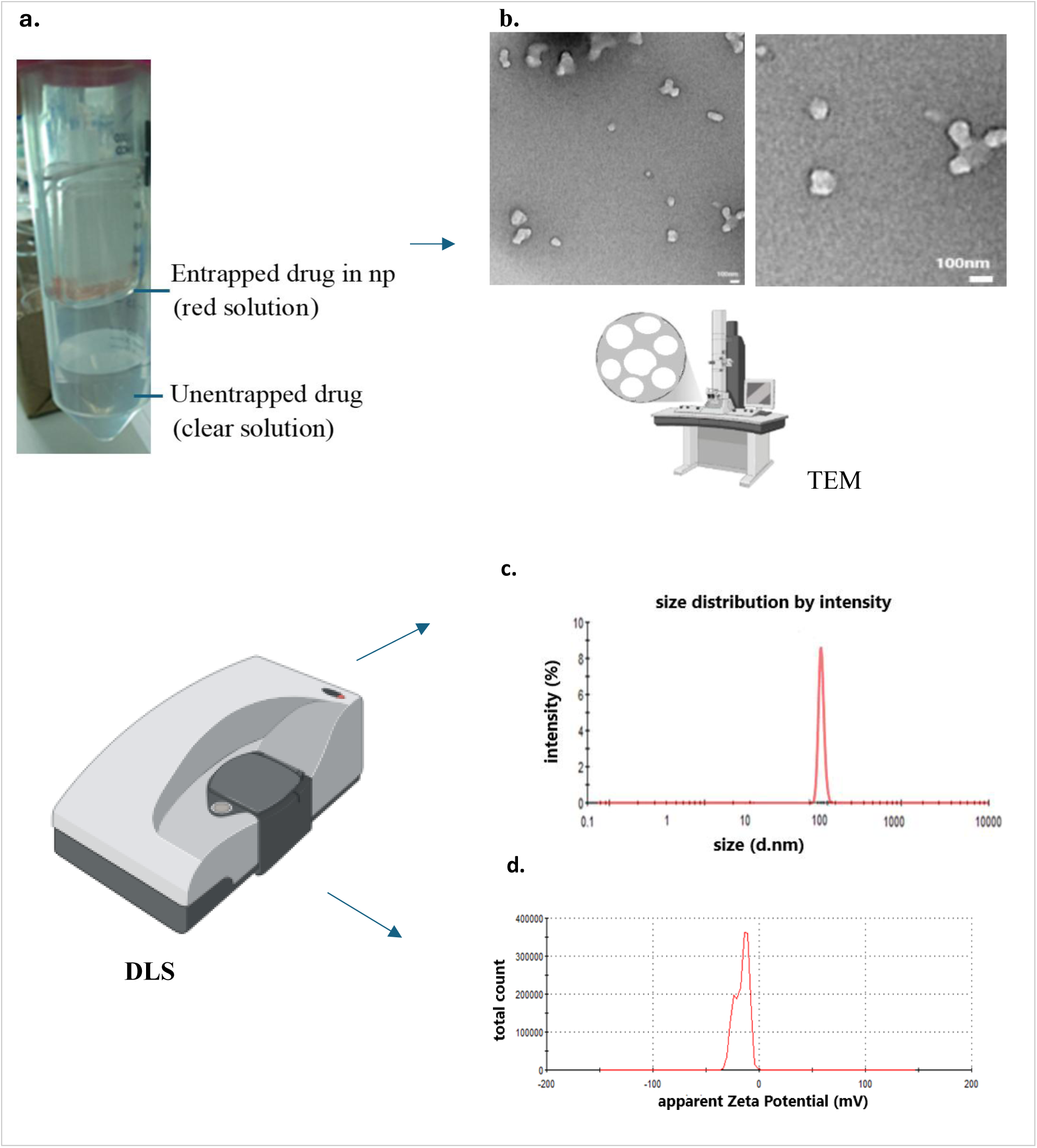
Nanoparticle characterization **a.** entrapped drugs (DOX) in nanoparticle (red solution) and unentrapped drugs (clear solution) separated by ultracentrifuge membrane filter tube (10 kDa MW pore size). **b** Representative TEM image of nanoparticle PLGA_PEG loaded Dox and Am21. **c** representative nanoparticle size distribution, **d** zeta potential of PLGA-PEG-A6 loaded DOX.AM21 (NPA6DOX.AM21)

**Table 2.**
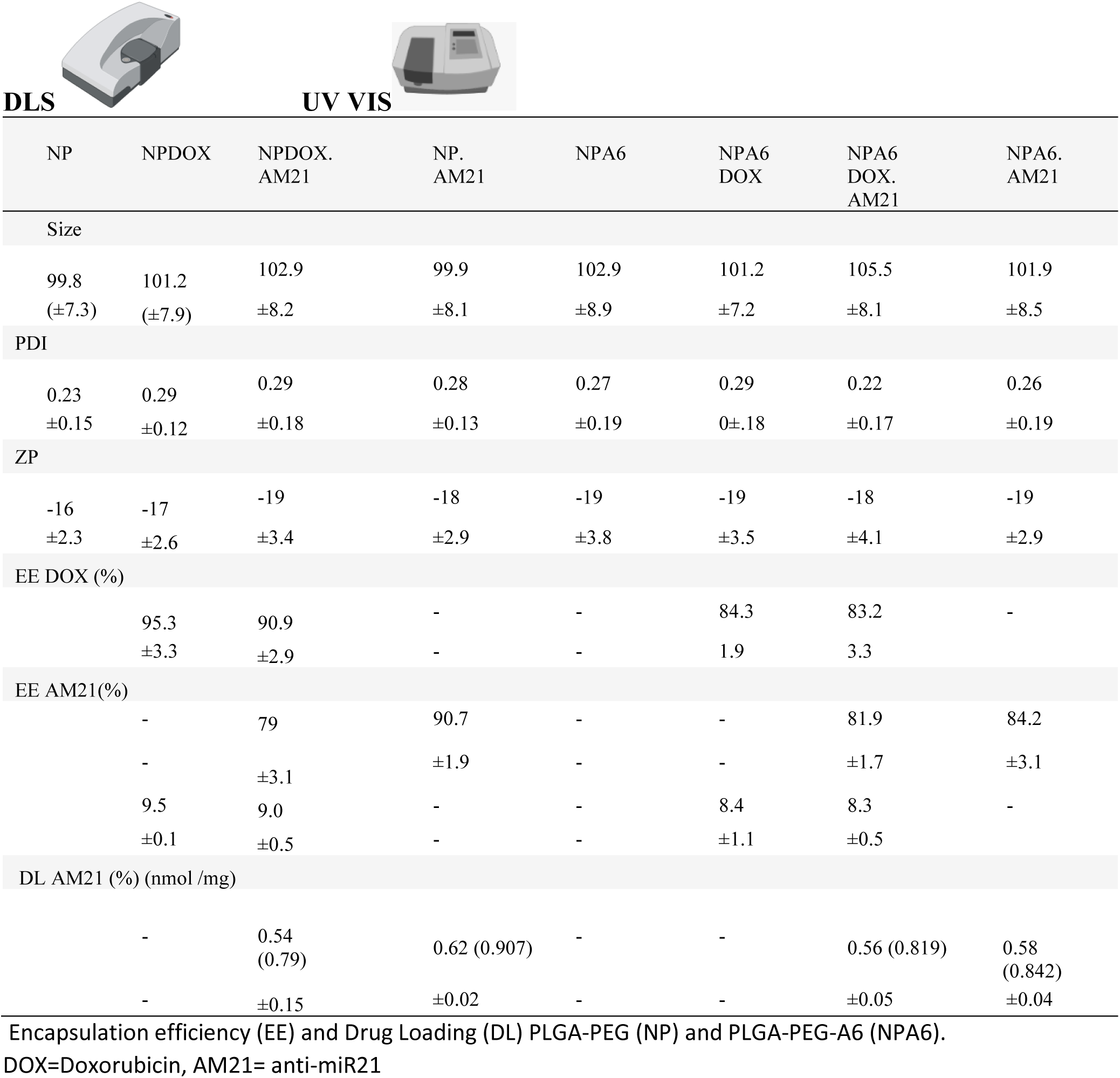
Size, PDI and ZP Encapsulation efficiency (EE) and Drug Loading (DL) of nanoparticles. PLGA(NP) and PLGA-A6 (NPA6)

| DLS |  | UV VIS |  |  |  |  |  |
| --- | --- | --- | --- | --- | --- | --- | --- |
| NP | NPDOX | NPDOX.<br>AM21 | NP.<br>AM21 | NPA6 | NPA6<br>DOX | NPA6<br>DOX.<br>AM21 | NPA6.<br>AM21 |
| Size |  |  |  |  |  |  |  |
| 99.8<br>($\pm 7.3$) | 101.2<br>($\pm 7.9$) | 102.9<br>$\pm 8.2$ | 99.9<br>$\pm 8.1$ | 102.9<br>$\pm 8.9$ | 101.2<br>$\pm 7.2$ | 105.5<br>$\pm 8.1$ | 101.9<br>$\pm 8.5$ |
| PDI |  |  |  |  |  |  |  |
| 0.23<br>$\pm 0.15$ | 0.29<br>$\pm 0.12$ | 0.29<br>$\pm 0.18$ | 0.28<br>$\pm 0.13$ | 0.27<br>$\pm 0.19$ | 0.29<br>$\pm 0.18$ | 0.22<br>$\pm 0.17$ | 0.26<br>$\pm 0.19$ |
| ZP |  |  |  |  |  |  |  |
| -16<br>$\pm 2.3$ | -17<br>$\pm 2.6$ | -19<br>$\pm 3.4$ | -18<br>$\pm 2.9$ | -19<br>$\pm 3.8$ | -19<br>$\pm 3.5$ | -18<br>$\pm 4.1$ | -19<br>$\pm 2.9$ |
| EE DOX (%) |  |  |  |  |  |  |  |
| | 95.3<br>$\pm 3.3$ | 90.9<br>$\pm 2.9$ | -<br>- | -<br>- | 84.3<br>1.9 | 83.2<br>3.3 | - |
| EE AM21(%) |  |  |  |  |  |  |  |
|  | - | 79 | 90.7 | - | - | 81.9 | 84.2 |
| | - | $\pm 3.1$ | $\pm 1.9$ | - | - | $\pm 1.7$ | $\pm 3.1$ |
| | 9.5<br>$\pm 0.1$ | 9.0<br>$\pm 0.5$ | -<br>- | -<br>- | 8.4<br>$\pm 1.1$ | 8.3<br>$\pm 0.5$ | - |
| DL AM21 (%) (nmol /mg) |  |  |  |  |  |  |  |
|  | - | 0.54<br>(0.79) | 0.62 (0.907) | - | - | 0.56 (0.819) | 0.58<br>(0.842) |
| | - | $\pm 0.15$ | $\pm 0.02$ | - | - | $\pm 0.05$ | $\pm 0.04$ |

Encapsulation efficiency (EE) and Drug Loading (DL) PLGA-PEG (NP) and PLGA-PEG-A6 (NPA6).
DOX=Doxorubicin, AM21= anti-miR21

The nanoparticles exhibited a spherical morphology under TEM (Fig 3). The negatively charged surface (zeta potential) of the nanoparticles makes them suitable for drug delivery as they can electrostatically bind to positively charged components on cell surfaces, facilitating drug transport across cell membranes [36]. Note that, the optimized formulation protocol presented in this study follows a series of formulation optimizations.

### 3.2 The Effect of formulation in vitro

#### 3.2.1 Evaluation of free Doxorubicin IC50 for resistant versus non-resistant cells

The IC_50_ of free doxorubicin in MDA-MB-231 resistant cells were found to be significantly higher, about 5 times higher compared to non-resistant MDA-MB-231 cells, measuring 25.65 µM and 4.559 µM, respectively (p<0.001) as shown in Fig. 4.1a The dose-response curve of the resistant cells shifted to the right, indicating a higher concentration of doxorubicin was needed to inhibit cancer cell growth, thus confirming drug resistance.

#### 3.2.2 Evaluation of nanoparticle formulation towards resistant MDA-MB-231/R cells

Comparison of mean cell viability across treatment showed that, the combination of DOX and AM21 loaded in NPA6 (NPA6DOX.AM21) exhibited the lowest cell viability, followed by NPA6DOX, NPA6AM21, NPDOX.AM21, NPDOX, and NP.AM21 respectively, in comparison to the free DOX drug and free AM21 Additionally, NPA6DOX.AM21 significantly inhibited cell growth more effectively than NPA6 Dox (Dox loaded alone) (p=0.01) and NPA6 AM21 (p=0.03). While NPDOX.AM21 displayed better inhibition than NPDOX (p=0.01) and) and NPAM21 (p=0.003) (Fig. 4.1b).

**Fig. 4.1.**
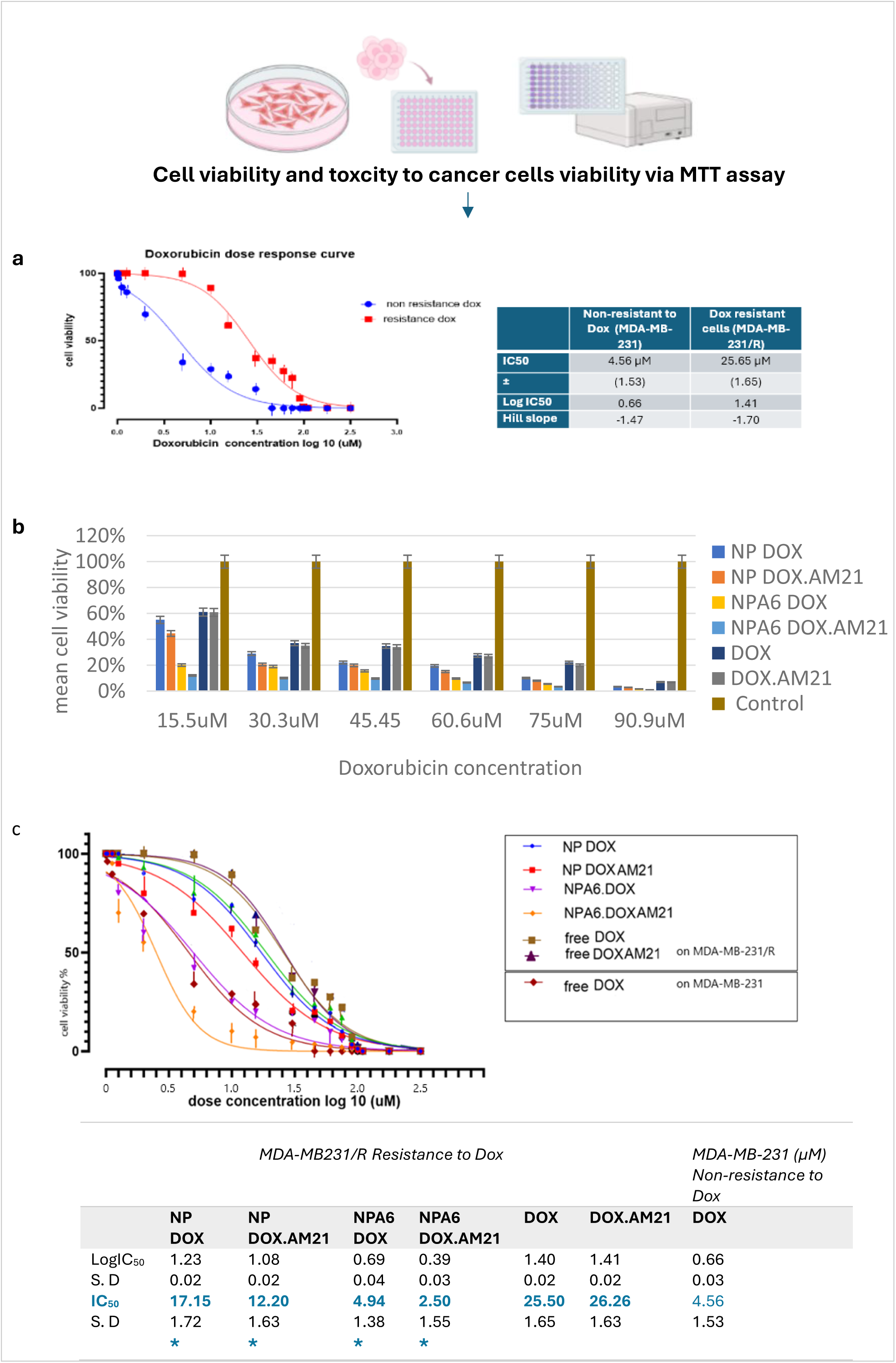
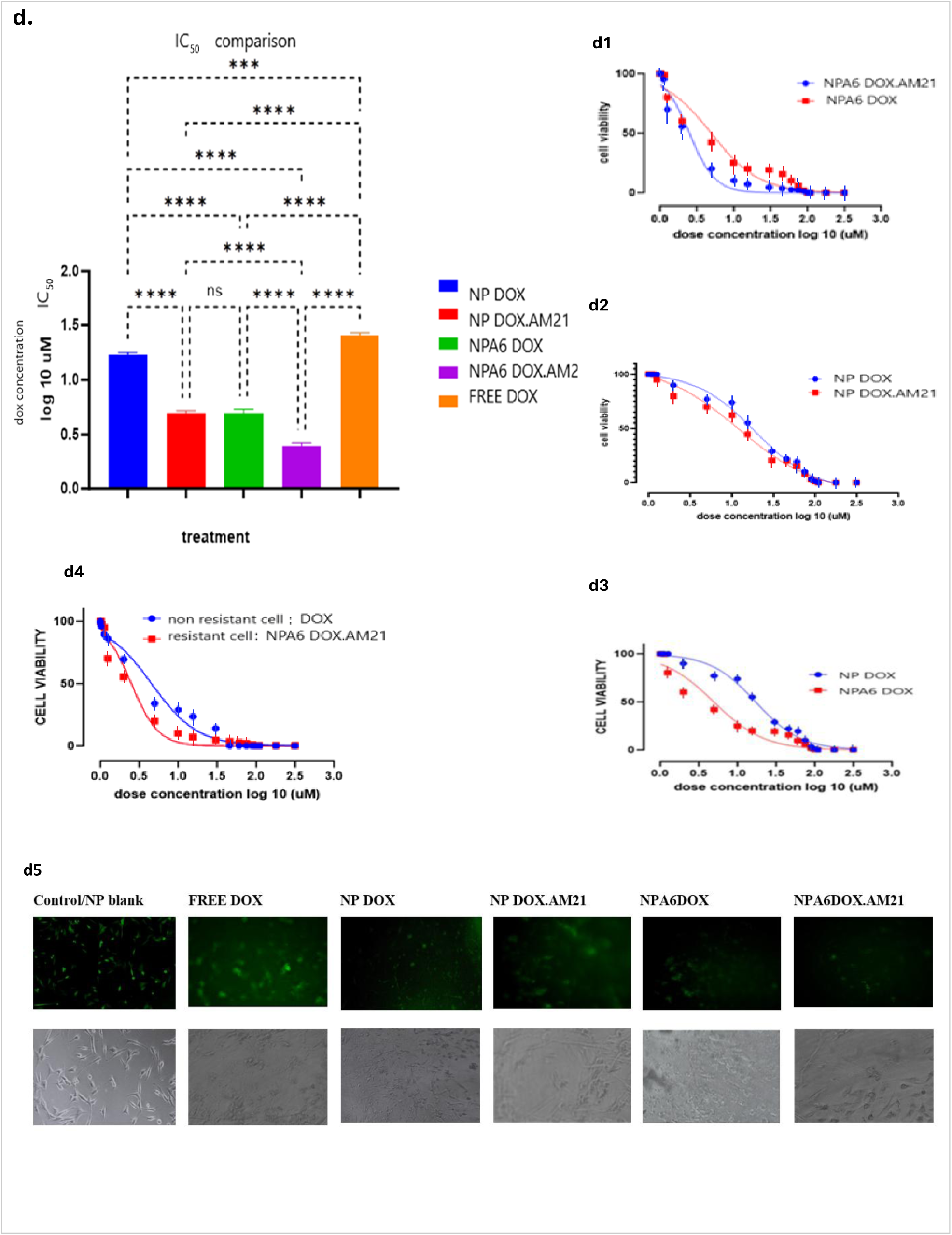
Assessment on drug resistant by cell viability and toxicity IC_50_. **a.** Comparative dose response curve and Dox IC50 (μM): between non-resistant to Dox (MDA-MB-231) cell and Dox-resistant (MDA-MB-231/R) cell both treated with free dox (p < 0.001) **b.** Mean cell viability of MDA-MB-231/R after 24 h exposure of treatments in each concentration in each treatment (except Negative control/ Blank NP). All NP and NPA6 loaded Dox had significant lower cell viability than the free doxorubicin at each concentration p < 0.01 (multi comparative ANOVA, post-hoc Bon Feroni) **c.** Comparison dose response curve and IC50 value all treatments (*p<0.01 significant difference of IC50 in comparison to free DOX in resistant MD-MB-231/R cell) **d.** Comparison dose response curve and IC50 value in resistant MDA-MB-231 cell (MDA/R). *P <0.001, ns=not significant (statistical ANOVA followed by Bon Ferroni). **d1, d2, d3:** comparison between np treatments in resistant MDA-MB-231 cell (MDA/R). d4: comparison between free Dox (tested on non-resistant cells) to NPA6DOX.AM21 (tested on resistant cells MDA/R) (independent comparison). **d5.** Representative MDA-MB-231/R cell viability under fluorescence microscopic after exposure to treatment in comparison to control. Green fluorescence (egfp) indicates cell viability. Low fluorescence /low cell viability of NP treatment compared to control.

To further investigate the impact of nanoparticles on cell resistance, IC_50_ values were compared across all nanoparticle formulations. IC_50_ was determined from the dose-response curve of logarithmic inhibitor concentration of Doxorubicin in micromolar (µM) to cell viability. Increasing doxorubicin concentration generally led to a decrease in cell viability across all treatments in resistant cells, (Fig. 4.1c). Both NP and NPA6 formulations exhibited a significant difference compared to free unencapsulated doxorubicin (p<0.01) (Fig. 4.1c, Fig. 4.1d), aligning with previous findings showing a lower doxorubicin dose in PLGA-loaded nanoparticles compared to the free form [37]. Notably, the lowest IC_50_ (was observed with NPA6.DOX.AM21 treatment, followed by NPA6DOX, NPA6 AM21, NPDOX AM21, NPDOX, DOX (p<0.01), highlighting the enhanced efficacy of killingd cancer cells with lower doxorubicin doses when incorporating the A6 peptide and AM21 in the nanoparticle cargo. Co-loading doxorubicin with AM21 resulted in a significantly lower IC_50_ compared to doxorubicin alone. For instance, IC_50_ values for NPA6DOX.AM21 versus NPA6DOX were 2.36 µM versus 4.94 µM respectively (p<0.001), and NPDOX.AM21 versus NPDOX were 12.2 µM versus 17.15 µM respectively (p<0.001) (Fig. 4.1c, Fig. 4.1d). This suggests that the incorporation of anti-miR21 into the DOX-Loaded nanoparticle formulation enhances the inhibition of cancer cells compared to DOX alone, while the A6 formulation combined with both loaded anti-cancer agents (DOX and AM21) significantly improved the effects on killing the resistant cancer cells. This similar enhanced effect due to treatment combination of Dox with AM21 were also previously observed in MCF-7 line [14] thus indicating, now it can be used for MDA-MB231 resistance cell. Additionally, A6 formulations demonstrated significantly lower IC_50_ values compared to non-A6 formulations (NPA6Dox versus NPDOX IC_50_ values were 4.94 µM versus 17.15 µM respectively, p<0.001, (Fig. 4c) while IC_50_ NPA6DOX.AM21 was lower than NPDOX.AM21 (P<0.01) (Fig. 4.1c, Fig. 4.1d). Targeted therapy (NPA6) in this study needed an extremely low dose of anticancer agents, while performing a better efficacy than the non-targeted. This may be due to its wider therapeutic windows and the flatten dose response curve [38] [39].

Finally, when comparing resistant cells treated with NPA6DOX.AM21 to non-resistant cells treated with free Dox (IC50 2.50 µM versus 4.55 µM respectively, p<0.001), the NPA6DOX.AM21 formulation exhibited an enhanced effect on resistant cells, potentially resensitizing these cells to treatment with almost 2x times lower doxorubicin dose.

Understanding the IC_50_ of Doxorubicin in MDA-MB-231 cells is crucial for patient care. The IC_50_, representing the dosage inhibiting 50% of cell growth, plays a vital role in assessing drug potency against cancer cells. This information may impact patient care by guiding treatment decisions.

Oncologists can tailor treatment based on IC_50_ values, exploring alternative therapies for patients with high IC_50_ indicating resistance. Moreover, personalized medicinal treatment can be achieved through individualized dosing strategies informed by IC_50_ levels, allowing optimization of drug efficacy and reduction of side effects based on patient responses. Furthermore, IC_50_ data informs research efforts to overcome drug resistance mechanisms [40], leading to the development of more effective therapies.. The current findings on IC_50_ in the MDA-MB-231 TNBC model indicate lower dosage Dox while co-loaded with anti-miR21 in targeted NPA6. This may translationally benefit lower risk off-targets toxicity as well as can serve as a reference for future research and applications.

#### 3.2.3 Assessing the miR-21 level and its target mRNA PTEN PDCD4 level after treatment with nanoparticle formulations

##### Resistance vs non-resistance

After plasmid transfection with the resistance gene into MDA-MB-231 cells, the expression level of miR-21 as well as PTEN and PDCD4 mRNA were assessed. Firstly, relative gene expression of resistant cells of MDA-MB-231/R was compared to the parental cell (MDA-MB-231non-resistant cell) **(Fig. 4.2)**. Using the Livak method the relative expression of the gene was compared to control by setting the control (non-resistant MDA-MB-231 cell) gene expression as 1. From Figure 4.2, miR-21 expression level in resistant cells was raised slightly to 1.246-fold in comparison to non-resistant cells. This increase, even though statistically not significant (p>0.05), was parallel to a previous study that showed an increase of miR-21 is in conjunction with the ABCB1 expression. Conversely, the gene expression levels of PTEN and PDCD4 were significantly downregulated to 0.220 and 0.004 (p<0.01) respectively. Although ABCB1 may or may not have a direct interaction with miR-21 and PTEN or PDC4 in the same pathway in this study, most previous studies showed that in the upstream pattern, miR-21 expression affects the P13k/PTEN pathway that regulates the ABCB1 expression. However, there was also another study showing that ABCB1 expression will loopback towards miR-21 expression by another mechanism [67] Chen et al. (2023) conducted a study that demonstrated a reciprocal regulation between ABCB1 and miR-21 in triple-negative breast cancer cells. Their findings indicated that increased ABCB1 expression led to the upregulation of miR-21, which negatively affected tumour suppressor genes and contributed to a resistant phenotype. The study highlighted the complexity of the feedback loop and suggested that targeting miR-21 could disrupt this cycle, potentially reversing resistance mediated by ABCB1. Furthermore, other studies showed that elevated miR-21 levels are parallel to the increase in ABCB1 expression, contributing to multidrug resistance in MDA-MB-231 cells [68] [69] [70]. However, due to limitations of the scope of the study, this current work had not verified the exact mechanism, but rather focused and assessed on the effect of ABCB1 transfection on miR-21 and its target mRNA expression.

**Figure 4.2:**
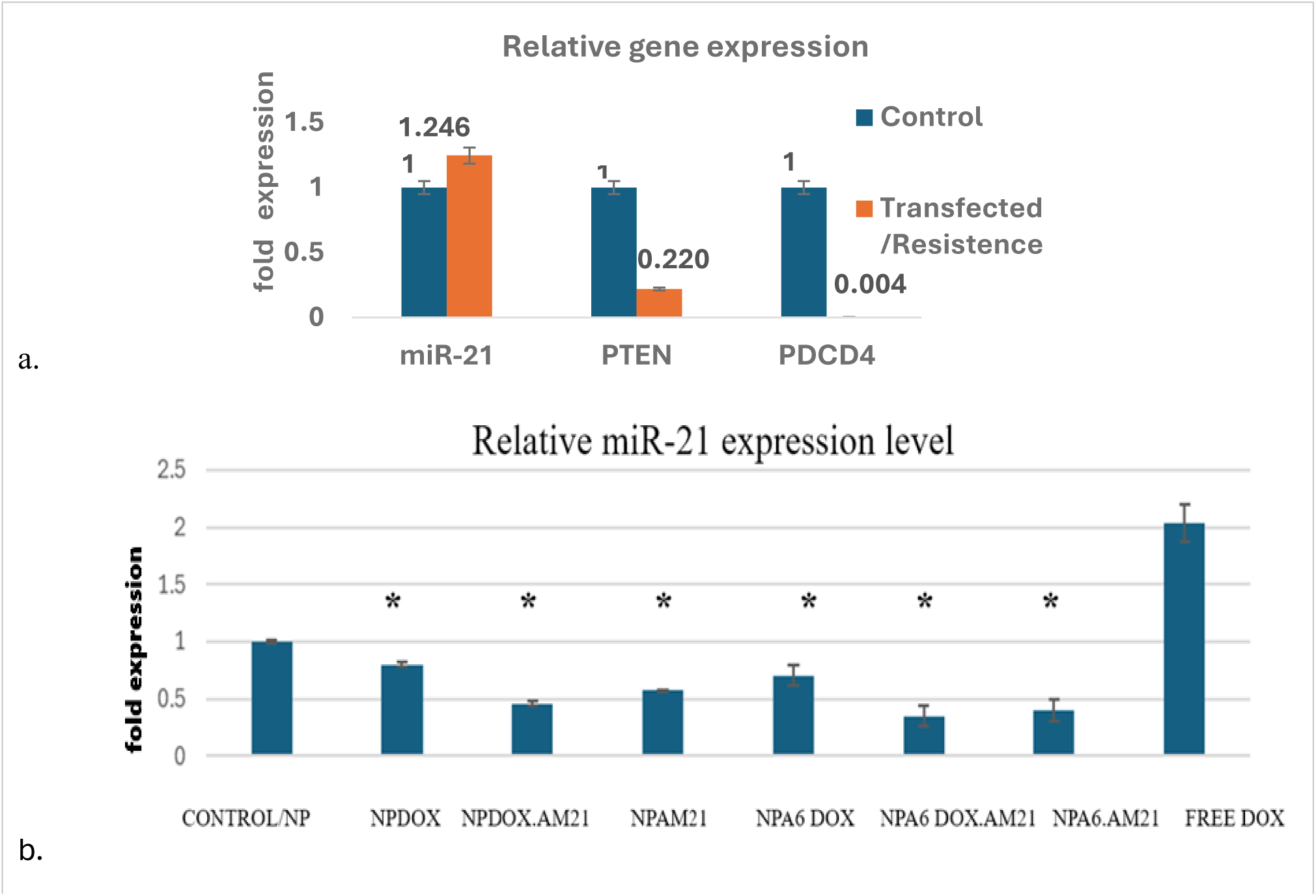
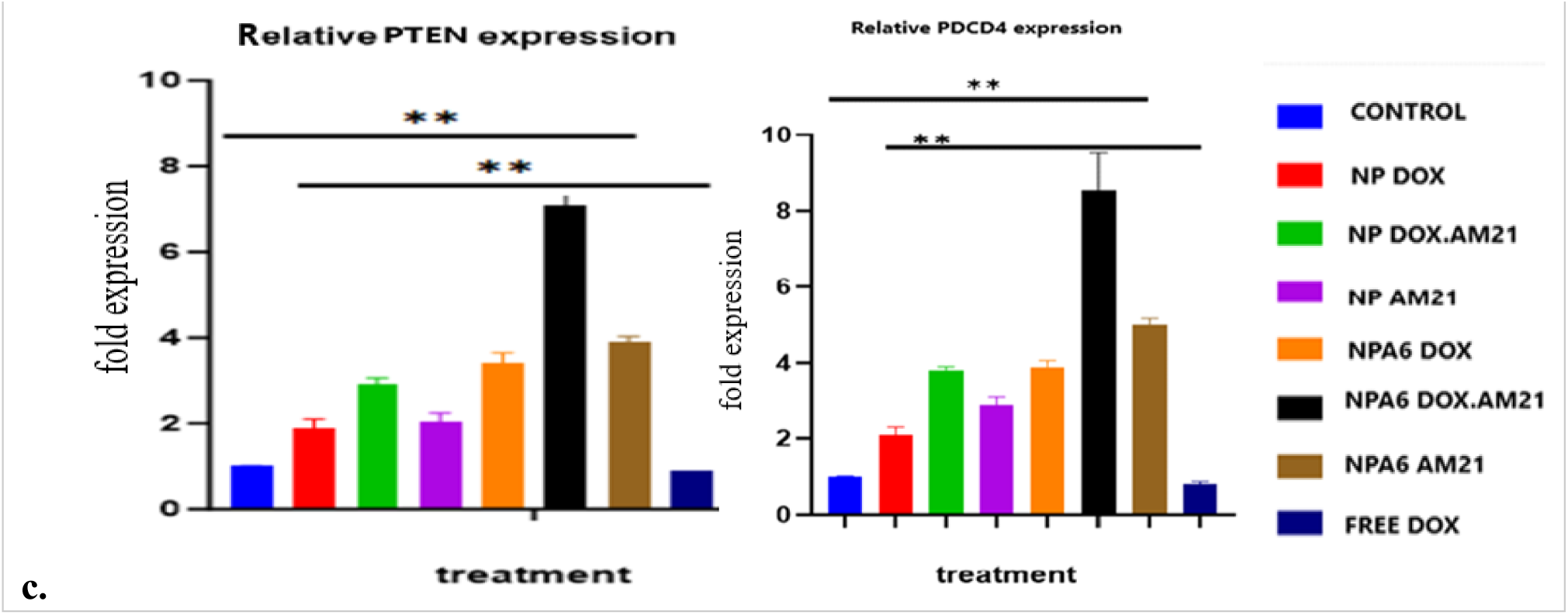
Relative expression of endogenous microRNA-21 and mRNA PTEN and mRNA PDCD4. **a.** Relative expression of resistance cell MDA-MB-231/R / (ABCB1 transfected) in comparison to non-resistance cell/non transfected) MDA-MB-231 (control). Control calibrated as 1. Less than 1 indicates downregulation, more than 1indicates upregulation **Downregulation** PTEN and PDCD4 mRNA expression level in MDA-MB-231/R (p< 0.01). miR-21 level was slightly increased in MDA-MB-231/R but was not statistically significant (p>0.01) student t-test. **b.** Relative miR-21 expression level across different treatments on the MDA-MB-231\R resistant cells. Control/ Blank NP (not containing drugs) was calibrated to 1 for relative gene expression, less than 1 indicates downregulation, more than 1indicates upregulation. NP and NPA6 formulations show a reduction of miR-21 expression, except free Dox relative to control, in resistant MDA-MB-231/R cells (*p<0.001 one way ANOVA, post hoc-Dunnets multi-comparison). **c.** Relative gene expression of PTEN and PDCD4 across treatment on the MDA-MB-231\R resistance cells. Blank NP (control) was set calibrated to 1 for relative gene expression. Less than 1 indicates downregulation, more than 1indicates upregulation. All nanoparticles (NP and NPA6 formulations) showed higher expression of (A) PTEN and (B) PDCD4 mRNAs compared to control and free Dox. *p<0.001 one way ANOVA, posthoc-Dunnets multi-comparison.

###### Testing nanoparticle formulation on the resistance cells

Following the finding established in the resistant cells, the nanoparticle formulation was tested on the resistance cells. Comparison of miR-21 level and mRNA PTEN and PDCD4 was assessed to see if there is significant difference between NPs.

###### Expression level of miR-21

The expression level of miRNA-21 was internally referred to as the housekeeping gene RNU6 while the expression level of the mRNAs was referred to the housekeeping gene beta-actin. For relative gene expressions, the control (NP/Blank) gene expression was calibrated as 1 as a standard. Figure 4.e2 shows the relative miR-21 level across different treatments compared to control/NP blank in the resistant cells. Treatment with free doxorubicin showed a 2-fold expression of miR-21 and this was expected as free doxorubicin was already being reported to be highly related to miR-21 expression in resistant cells. However, the nanoparticle formulation showed reduction of miR-21 with the low 2-fold reduction in NPA6 DOX.AM21 treatment followed by NPA6.AM21, NP DOX.AM21, NP.AM21, NPA6DOX, AND NP DOX respectively, relative to control. All NP groups loaded with AM21 showed less miR-21 expressions relative to control (Blank NP). Dox loaded NP and NPA6 without AM21 slightly reduced the miR-21 expression, relative to control indicating underlying mechanism that Dox loaded NP system may partly play a role in miR-21 expression intracellularly. The PLGA-PEG has been known to facilitate the entry of drugs into the cells, and will biodegrade in the cells and released its contents [24]. Doxorubicin is an anti-cancer agent and is toxic to DNA. Once it is inside the cell’s nucleus it can intercalate with DNA to stop cell replication and proliferation [66]. Furthermore, Dox can also generate reactive oxidative species (ROS) and could damage the intracellular proteins or molecules. This doxorubicin-DNA intercalation and ROS process may interfere with the rest of any gene replication including miR-21 gene, as DNA being damaged by DOX, causing cell apoptosis and inhibition to any gene transcription and translation. And this could be presented by the slight reduction of endogenous miR-21 level in cells treated with NP loaded DOX, even without AM21. For free Dox, as previously described, entry into cells is mediated by membrane diffusion, thus a different mechanism may play a role in how free Dox induced the miR-21 level compared to NP loaded Dox. Furthermore, NP loaded DOX cellular entry was by endocytosis and can “escape’ the ABCB1 efflux transporter that may have exerted a different effect on miR-21 level. On another note, one previous study revealed that blank PLGA-PEG may minimally reduce the miR-21 level in the cells (but this reduction may be negligible as most PLGA-PEG NPs will be degraded in the cells) [24], thus the minimal decrease of miR-21 due to the NP loaded DOX (without AM21) was expected as in this current study. As miR-21 decreased, this led to an increased expression in its target genes mRNA, PTEN and PDCD4 as miR-21 normally functions to suppress PTEN and PDCD4. The endpoint of the result showed that NPA6 DOX AM21 had the highest reduction of miR-21 level compared to all treatments and to this control. This finding may greatly contribute to overcoming miR-21 related-drug resistance in chemotherapy.

###### PTEN and PDCD4 mRNA level

Multi-comparison between treatments showed that combination of doxorubicin with AM21 loaded in NP and NPA6 had higher PTEN, PDCD4 mRNA expression than single loaded dox or single loaded AM21. Figure 4.e3 shows the PTEN and PDCD4 expressions after treatment with nanoparticle formulations. The NPA6 DOX.AM21 formulation had the highest expression for PTEN and PDCDC4 followed by NPA6.AM21, NPA6DOX, NP DOX.AM21, NP.AM21 and NP DOX and free doxorubicin in comparison to control (blank NP). The NP or NPA6 formulations loaded with AM21 had antisense to miR-21, and thus hypothetically may inhibit the endogenous miR-21 resulting in a low miR-21 level. Hence, there would be less miR-21 available to bind to the downstream PTEN and PDCD4 mRNA, indirectly causing the elevation of PTEN and PDCD4 in the cells. Furthermore, in the formulation with the AM21 co-loaded with Dox, this could enhance the PTEN and PDCD4 expression as the genes are responsible for controlling cell proliferation in apoptosis and cell death pathway. This agrees with the cell viability assay and the IC_50_ results, in which NPA6 and NP loaded with AM21 and dox had the lowest IC_50_, suggesting its enhanced ability to reduce cell resistance.

This study confirms that the inhibition of miR-21 via anti-miR-21 leads to lower levels of miR-21 presented in the gene expression data. As miR-21 normally functions to regulate PTEN and PDCD4 genes by suppressing their expression, the inhibition of miR-21’s function via AM21 allowed the PTEN and PDCD4 mRNA to rise. These two genes PTEN and PDCD4 are involved in programmed cell death and apoptosis, hence by increasing their expression level, their original function to regulate the cell cycle and partly downregulate cell resistance could be restored. This restoration could help to re-sensitize the resistant cells to drug treatment. As the current work had presented, a lower dose of Doxorubicin (shown as low IC_50_) was needed to kill the cells when it was co loaded with anti-miR-21 (AM21).

### 3.3 Efficacy of Nanoparticle Formulations in a Mouse Tumor Model: Tumor Size Reduction

In this preclinical study, tumor size inhibition served as a primary indicator of nanoparticle treatment efficacy, as reduced size indicates cancer cell proliferation inhibition or tumor cell death induction. Assessing tumor size reduction in mouse models is critical for predicting clinical outcomes in human [41]

#### The study evaluated tumor size over 24 days post-initial drug injection in mice with an average body weight between 24g-32g at the study’s end. (Fig. 5)

On the 24th day, mean tumor volumes of treated groups were compared to controls and free doxorubicin, as well as multiple comparisons were performed between treatments (<u>Fig.5b</u>). In terms of overall tumor reduction from highest to lowest, the NPA6DOX.AM21 > NPA6DOX > NPDOX > free doxorubicin showed 54% > 45% > 33.5% >15% tumor size reduction respectively, in comparison to control (blank NP).The mean tumor volume for NPA6 loaded with doxorubicin and AM21 (NPA6DOX.AM21) exhibited the smallest size, indicating successful tumor inhibition and significantly reduced from both the control (p<0.001) and in comparison to the rest of other treatments. Notably, NPA6DOX.AM21 demonstrated 45%, 30% and 14.4% tumor reduction compared to free doxorubicin NPDOX and NPA6DOX groups, respectively (p<0.01).

**Fig. 5.**
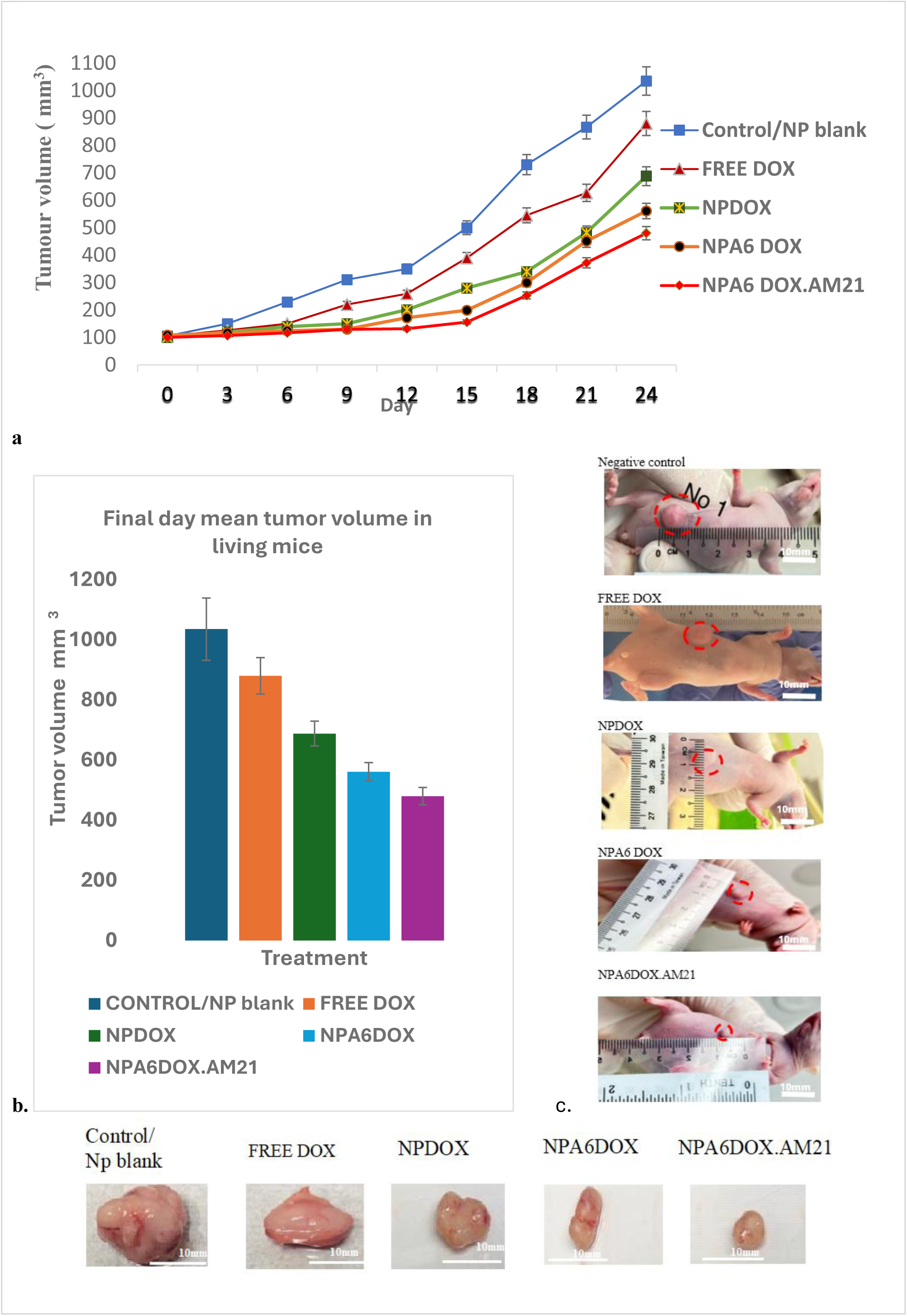
Assessment on tumor size. **a.** Tumor growth from 0 to 24 days after receiving different nano formulation treatment **b.** comparative mean tumor size at final day (day 24) **p<0.05 ***p<0.001 **c.** representative image of tumor (blue circle) in tumor bearing mice treated with the np formulations **d.** representative image of tumor ex vivo from the mice receiving treatments and control (white scale bar: 10mm.)

All nanoparticles (NPs) and NPA6 significantly inhibited tumor growth compared to the control (p<0.001) and free doxorubicin (p<0.01). NPA6 loaded drugs (NPA6 dox, NPA6 DOXAM21) exhibited enhanced inhibition (18.5% and 30.2% reduction) compared to NPDOX (p<0.05), indicating that the targeting peptide on NPA6’s surface improved inhibition of tumor growth. This study had anticipated substantial tumor shrinkage, suggesting the need for expand investigation in future clinical trials for the development of new cancer therapeutic.

### 3.4 The effects of the prepared formulation on other off targe**t organs**

#### 3.4.1 Relative Doxorubicin intensity in tumor

The study assessed the biodistribution of doxorubicin-loaded nanoparticle in mice’s tumor and vital organs to evaluate its specificity and safety. The fluorescence intensity of doxorubicin in the tumor was used to assess nanoparticle accumulation, with NPA6-loaded drugs (NPA6 DOX and NPA6DOX.AM21) showing significantly higher fluorescence compared to free doxorubicin and NPDOX. This suggests that the A6 peptide on the NP facilitated enhanced doxorubicin delivery and accumulation in the tumor compared to free doxorubicin and NP without the A6 peptide (illustrated in Figure 6).Statistical analyses supported these findings, indicating their significance (p<0.01 These results are consistent with previous studies showing that A6 targeting leads to inhibited tumor growth and improved survivability in tumor-bearing mice [42].

**Fig. 6.**
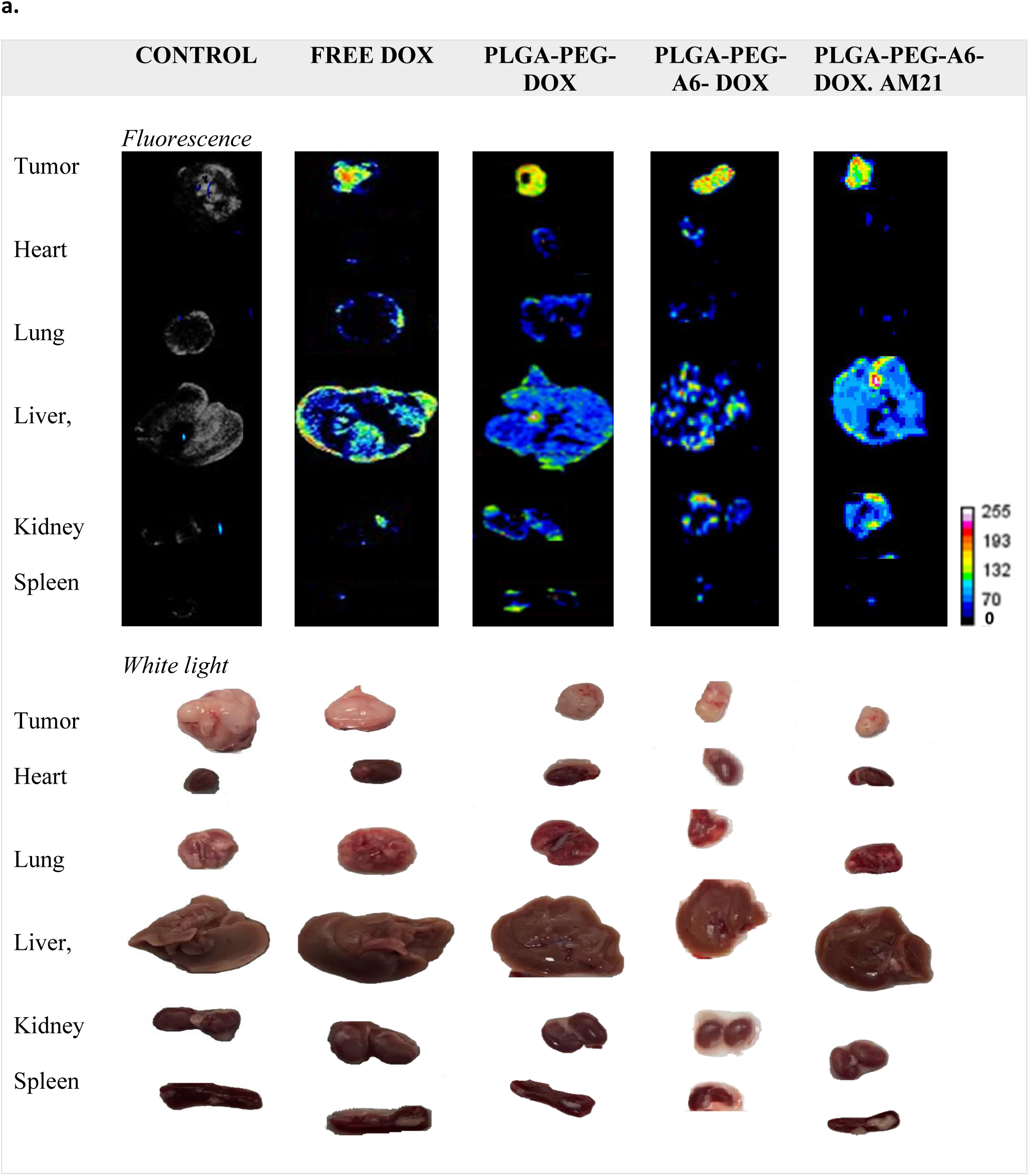
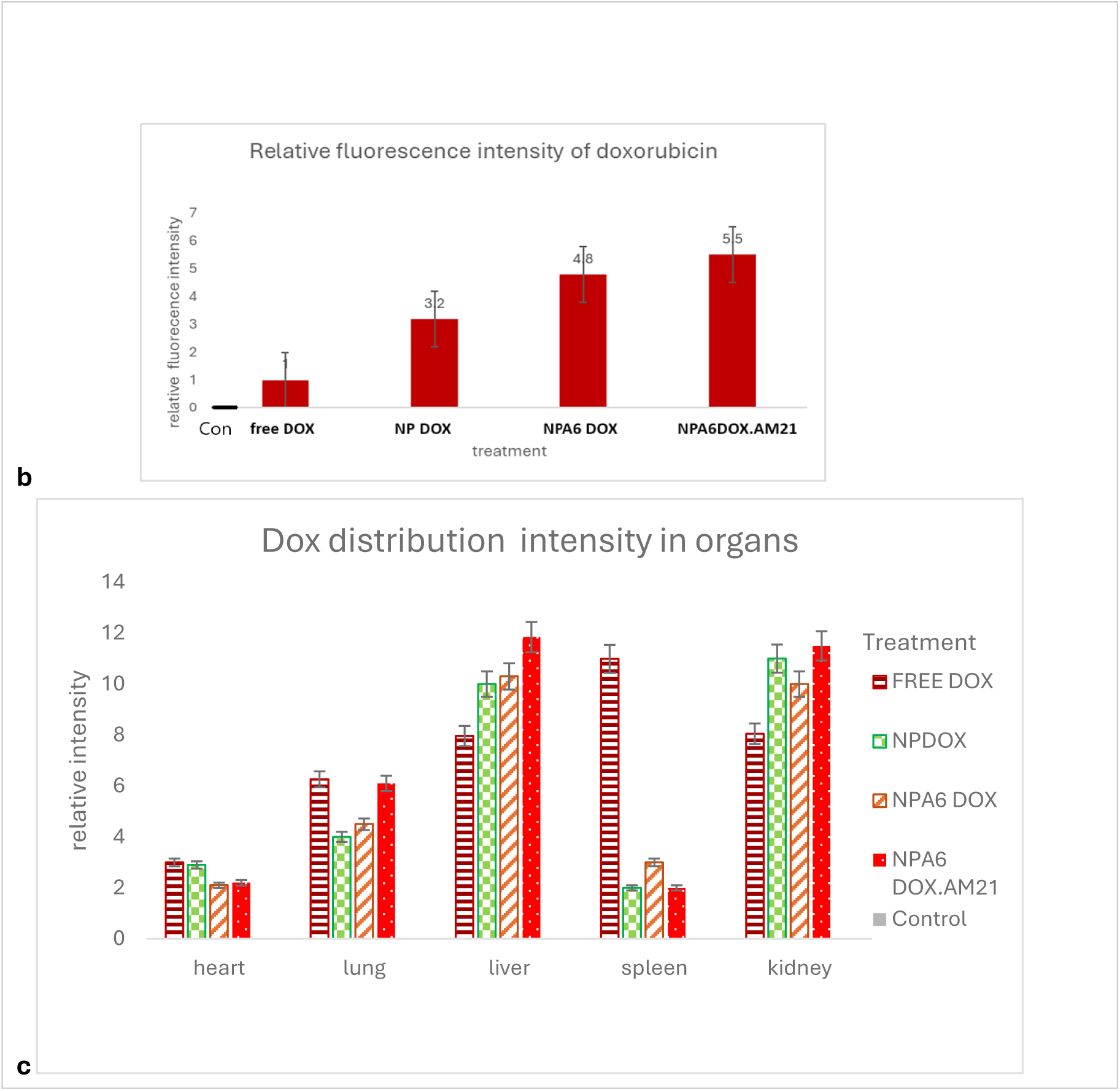
Ex vivo Dox fluorescence biodistribution after 24hr tail vein injection with treatments of nanoparticle drug and free doxorubicin (via FX Pro Bioimaging) **a.** Ex vivo image of tumours and organs with Doxorubicin signal fluorescence. **b** relative Dox fluorescence intensity in tumour. Highest fluorescence intensity indicating the accumulation of Dox in tumours with treatment of NPA6DOX AM21 followed by, NPA6Dox and NPDOX in comparison to free dox (p<0.05) while NPA post hoc. **c.** Relative dox fluorescence intensity in organs

The current study provided insights into the pharmacokinetics of a newly formulated drug. Typically, when a drug enters the human body, it undergoes four stages: absorption, distribution, metabolism, and excretion (ADME). Absorption involves the movement of the drug from the site of administration to the target site. For instance, following oral administration, absorption occurs in the small intestine before entering the bloodstream for distribution throughout the body. Distribution refers to the spread of the drug to various tissues and organs In previous doxorubicin studies, researchers often focus on its distribution in organs like the liver, kidney, lungs, spleen, and heart, besides the tumor site. [43]. Once the drug enters organs like the liver, the drug undergoes metabolism. The liver is crucial for drug metabolism, where the drug molecules are broken down. Subsequently, the drug is excreted, primarily through the kidneys. These stages play a significant role in determining the drug’s fate.[44]

In this study, mice were injected with nanoparticles containing the drug nanoparticle formulations via the tail vein, leading to direct entry into the distribution phase through the blood circulation. The study evaluated the distribution of the nanoparticles carrying the drugs to major organs. In blood vessels, the small size of the nanoparticles (100-200nm) has the enhanced permeability and retention (EPR) effect, in which these nanoparticles can efficiently enter tumor sites via leaky blood vessel membranes [45][31][46]. Moreover, the PEG coating on the PLGA surface increases circulation time, aiding in tumor accumulation and penetration [47] As a result, the nanoparticle-loaded drugs showed a higher fluorescence signal accumulation compared to free doxorubicin (p<0.01). By incorporating the A6 peptide targeting on the nanoparticles, there was an improved selective binding to TNBC cells, leading to enhanced fluorescence signals with NPA6 loaded drugs compared to non-A6 nanoparticles and free doxorubicin.

#### 3.4.2 Fluorescence intensity of Doxorubicin in organs

This study further assessed the distribution of doxorubicin-loaded nanoparticles in off-target organs by examining the relative doxorubicin fluorescence intensities. While there was no significant increase compared to free doxorubicin in the heart and lungs (p<0.05), a slight but not significant increase was detected in the kidneys and liver. The PEG coating helped in escaping opsonization from blood proteins in systemic circulation. [48]

This study highlighted the superior accumulation of NPA6-formulated drugs in tumors. (Fig. 6). Additionally, the benefits of nanoparticle accumulation could potentially be utilized for treating other cancer types, like liver cancer, due to the liver’s role in metabolizing larger molecules. Adjusting nanoparticle size may lead to liver accumulation, as previous studies have shown. [49][50] These advancements in nanotechnology have paved the way for drug development and clinical trials.

The study also noted that the fluorescence intensity of doxorubicin can be immediately captured after ex vivo excision of the tissue from the mice. Doxorubicin, when administered at high doses (such as 10 mg/kg), can be detected by fluorescence in ex vivo tumors and organs within 24 hours. However, our additional observation showed that after 48 and 72 hours, the fluorescence of doxorubicin cannot be detected, indicating rapid clearance in the blood circulation. Additional detection tools, such as blood plasma clearance analysis, were not utilized in this study but are necessary to understand the pharmacokinetics of doxorubicin in mice.[51]

This study aims to emphasize the effectiveness of the tumor-homing peptide strategy (A6 peptide). Homing peptides, which can bind to specific receptors on cell surfaces, are a targeted approach in cancer nano therapy [52] [53]. They enhance drug delivery to the tumor site, improving treatment efficacy while minimizing side effects.

Previous studies have shown that formulations including the A6 peptide have better anticancer effects than those without it, highlighting their potential in cancer therapy [54][55][56] Homing peptides is one of targeted approach in cancer nano-therapy as it can bind to specific receptors on the surface of cells [57] It enhances delivery of drugs to the tumor site, thus improving the treatment while reducing side effects[58][16] Furthermore the nanodevices’ physicochemical properties play a crucial role in their entry into cells and efficacy in targeting tumors [59][60]. As previously described, our nanoparticle characterization may fit the suitability due to its surface charges and np sizes, peg coating and peptide targeting.

Histological analysis of organs from mice treated with the nanoparticle formulation confirmed the safety and lack of toxicity in vital organs, particularly the heart (Fig. 7). Unlike free doxorubicin that can cause toxic effects on the heart, the nanoparticle formulation showed no significant structural changes or damage in heart tissues. Further toxicological studies are required to determine the optimal dosage of the nanoparticle formulation. Overall, the study suggests that low doses of doxorubicin in nanoparticle formulations effectively target and kill cancer cells while minimizing systemic toxicity.

**Fig. 7.**
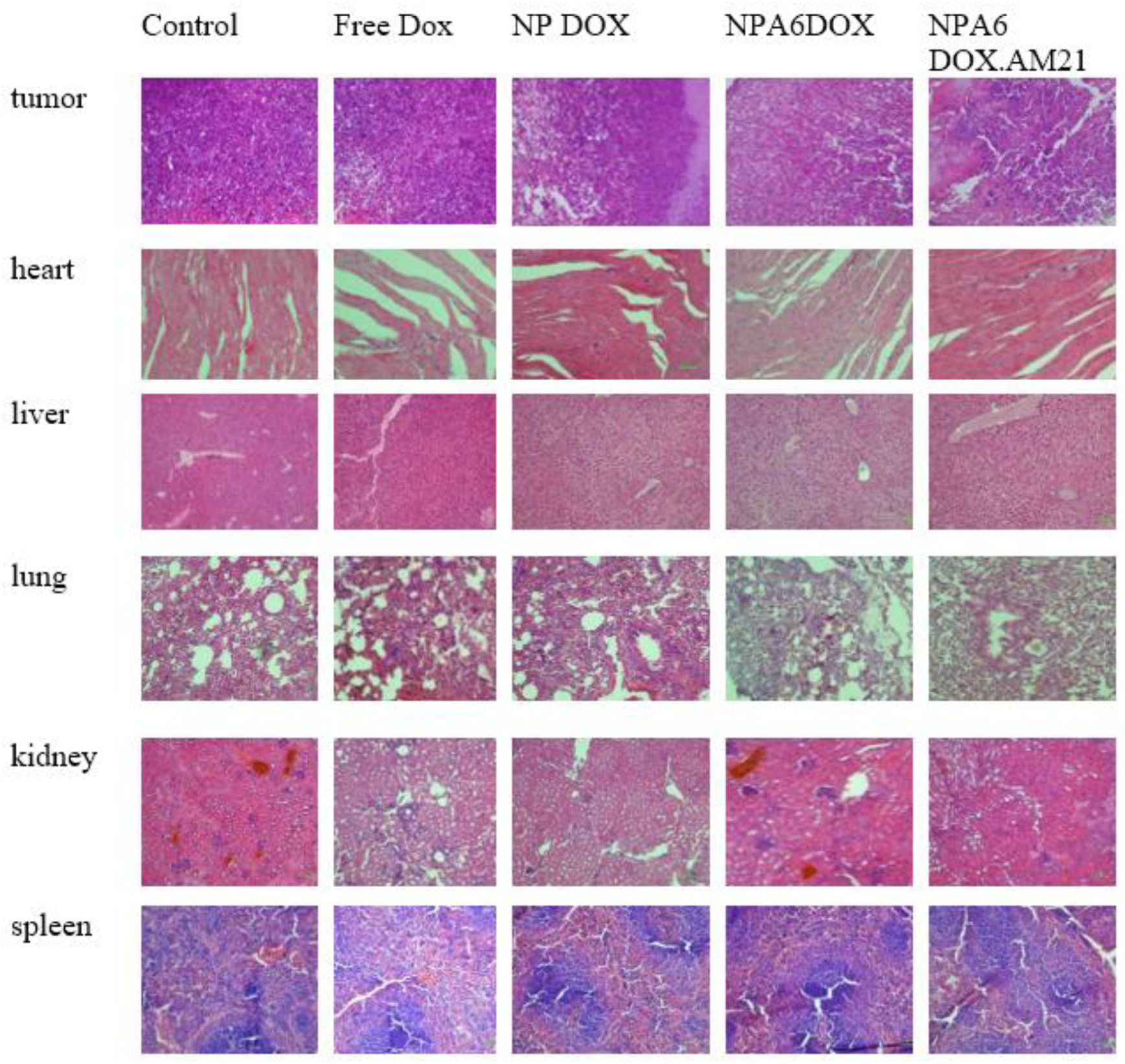
Image of histology (stained with H&E) of tumor and organs from mice treated with DOX and np formulation (under 200X magnification). There is no significant toxicity in tissues in mice treated with np formulation in off target organs

#### Toxicity Analysis and Future Research Directions

In our in vitro efficacy study result, it is noted that low dosage of Dox in NP formulation is sufficient in effectively killing the cancer cells. This may positively translate into reduction of the risk of systemic toxicity dissemination in healthy tissues and in metabolic and excretion organs [61]. Our NP formulation showed no toxic effect on the organs, however pharmacokinetic study and future research based on toxicity analysis is crucial, especially considering the potent and toxic nature of free doxorubicin towards healthy cells [62]. Future toxicity studies should focus on determining the lowest doxorubicin concentration [63]required to effectively target tumors while minimizing the risk of organ failure in systemic model.

This study highlights the successful development of PLGA-PEG-A6 nanoparticles loaded with doxorubicin and antisense miR-21 using modified double emulsion solvent evaporation. The incorporation of the targeting A6 peptide on the PLGA-PEG nanocarrier improved drug delivery efficiency and anticancer efficacy. Notably, this study represents the first instance of combining PLGA-PEG-A6 peptide with doxorubicin and antisense miR-21, aiming to address multidrug resistance in TNBC cells and enhance specific targeting in vivo. The co-loading of antisense miR-21 with doxorubicin in the NPA6 nanocarrier demonstrated a reduction in drug resistance and increased sensitivity to doxorubicin in TNBC cells with low dosage of DOX. This potentially offers translational benefits by minimizing systemic toxicity to healthy cells. The use of peptide and antisense microRNA technology in cancer therapeutics [64] [65] shows promise in overcoming drug resistance and enhancing cancer cell inhibition, paving the way for nucleic acid-based therapies to combat cancer effectively.

## Conclusion

The novel formulation of PLGA-PEG conjugated with A6 peptide loaded with combination doxorubicin and anti-miR-21, enhanced toxic effects on drug resistant-cell of MDA-MB-231 and re-sensitized the cell to doxorubicin treatment in vitro. Furthermore, the PLGA-PEG-A6 peptide loaded drugs targeted and reduced tumor size while diminished off target accumulation in organs in mice model offering a safety treatment.

In conclusion, the formulation developed in this study represents a promising advancement in the targeted treatment of triple negative breast cancer and thus holds great potential for an effective, better-tolerated therapeutics outcome for patients

## Declaration

### Ethics

This research had been approved by Animal ethics research board from the Institutional Animal Care and Use Committee (IACUC), Faculty of Medicine, University Malaya (UM) MALAYSIA (2023-240105/PHARM/R/HA).

### Consent for publication

Not applicable

### Availability of data and materials

Supplementary Information in (S.Info) file.

### Authors’ contributions

Z.I: writing reviewing, editing the original draft, project conceptualization, experimental investigation, created the nanoparticle and tested in vitro and in vivo, data curation, data analysis and visualization, funding acquisition through proposal writing.

CPP: main project supervision, reviewing and editing manuscript, funding acquisition

ZC: reviewing and editing manuscript, project supervision, funding acquisition

HA: contributed partly to experimental work (methodology in nanoparticle development and in vivo work)

MA: contributed to histology analysis

## Acknowledgements

The authors appreciate the support of the instrumentation and laboratory facilities in Taylors University and University of Malaya that is essential in vitro and in vivo experimentation. The author also acknowledges a working colleague Dr. Sareh Kamran and all colleague and laboratory staff at UBAT laboratory at University of Malaya and Research Lab 3, School of Bioscience at Taylors University, Malaysia.

## References

1. H. A. Wahba and H. A. El-Hadaad, Current approaches in treatment of triple-negative breast cancer. Cancer Biol. Med. 12, 106–116 (2015). 10.7497/j.issn.2095-3941.2015.0030

2. I. C. Salaroglio, E. Gazzano, A. Abdullrahman, E. Mungo, B. Castella, et al., Increasing intratumor C/EBP-β LIP and nitric oxide levels overcome resistance to doxorubicin in triple negative breast cancer. J. Exp. Clin. Cancer Res. 37, 1–20 (2018). 10.1186/s13046-018-0967-0

3. A. N. Linders, I. B. Dias, T. López Fernández, C. G. Tocchetti, N. Bomer, et al., A review of the pathophysiological mechanisms of doxorubicin-induced cardiotoxicity and aging. Npj Aging 10, (2024). 10.1038/s41514-024-00135-7

4. D. M. Brown, C. Croce, and S. P. Nana-Sinkam, Chapter 2 - Clinical and Therapeutic Applications of MicroRNA in Cancer. in Transl. MicroRNAs to Clin., edited by J. Laurence (Academic Press, Boston, 2017), pp. 17–37

5. Y. H. Feng and C. J. Tsao, Emerging role of microRNA-21 in cancer (Review). Biomed. Reports 5, 395–402 (2016). 10.3892/br.2016.747

6. Y. Zhang, B. Xu, and X. P. Zhang, Effects of miRNAs on functions of breast cancer stem cells and treatment of breast cancer. Onco. Targets. Ther. 11, 4263–4270 (2018). 10.2147/OTT.S165156

7. J. Chen, C. Zhou, J. Li, X. Xiang, L. Zhang, et al., MiR-21-5p confers doxorubicin resistance in gastric cancer cells by targeting PTEN and TIMP3. Int. J. Mol. Med. 41, 1855–1866 (2018). 10.3892/ijmm.2018.3405

8. S. Zhu, H. Wu, F. Wu, D. Nie, S. Sheng, et al., MicroRNA-21 targets tumor suppressor genes in invasion and metastasis. Cell Res. 18, 350–359 (2008). 10.1038/cr.2008.24

9. Z. X. Wang, B. Bin Lu, H. Wang, Z. X. Cheng, and Y. M. Yin, MicroRNA-21 Modulates Chemosensitivity of Breast Cancer Cells to Doxorubicin by Targeting PTEN. Arch. Med. Res. 42, 281–290 (2011). 10.1016/j.arcmed.2011.06.008

10. S. Raniolo, V. Unida, G. Vindigni, C. Stolfi, F. Iacovelli, et al., Combined and selective miR-21 silencing and doxorubicin delivery in cancer cells using tailored DNA nanostructures. Cell Death Dis. 12, 3–11 (2021). 10.1038/s41419-020-03339-3

11. G. Da Yang, T. J. Huang, L. X. Peng, C. F. Yang, R. Y. Liu, et al., Epstein-Barr Virus_Encoded LMP1 Upregulates MicroRNA-21 to Promote the Resistance of Nasopharyngeal Carcinoma Cells to Cisplatin-Induced Apoptosis by Suppressing PDCD4 and Fas-L. PLoS One 8, 1–15 (2013). 10.1371/journal.pone.0078355

12. X. Wei, W. Wang, L. Wang, Y. Zhang, X. Zhang, et al., MicroRNA-21 induces 5-fluorouracil resistance in human pancreatic cancer cells by regulating PTEN and PDCD4. Cancer Med. 5, 693–702 (2016). 10.1002/cam4.626

13. L. Hong, Y. Han, Y. Zhang, H. Zhang, Q. Zhao, et al., MicroRNA-21: a therapeutic target for reversing drug resistance in cancer. Expert Opin. Ther. Targets 17, 1073–1080 (2013). 10.1517/14728222.2013.819853

14. M. Rui, Y. Qu, T. Gao, Y. Ge, C. Feng, et al., Simultaneous delivery of anti-miR21 with doxorubicin prodrug by mimetic lipoprotein nanoparticles for synergistic effect against drug resistance in cancer cells. Int. J. Nanomedicine 12, 217–237 (2017). 10.2147/IJN.S122171

15. J. V. McGowan, R. Chung, A. Maulik, I. Piotrowska, J. M. Walker, et al., Anthracycline Chemotherapy and Cardiotoxicity. Cardiovasc. Drugs Ther. 31, 63–75 (2017). 10.1007/s10557-016-6711-0

16. J. Li, Q. Wang, G. Xia, N. Adilijiang, Y. Li, et al., Recent Advances in Targeted Drug Delivery Strategy for Enhancing Oncotherapy. Pharmaceutics 15, (2023). 10.3390/pharmaceutics15092233

17. A. Sharma, N. Jain, and R. Sareen, Nanocarriers for diagnosis and targeting of breast cancer. Biomed Res. Int. 2013, 960821 (2013). 10.1155/2013/960821

18. H. A. Ebrahimi, Y. Javadzadeh, M. Hamidi, and M. Barzegar Jalali, Development and characterization of a novel lipohydrogel nanocarrier: Repaglinide as a lipophilic model drug. J. Pharm. Pharmacol. 68, 450–458 (2016). 10.1111/jphp.12537

19. R. Devulapally, N. M. Sekar, T. V Sekar, K. Foygel, T. F. Massoud, et al., Polymer Nanoparticles Mediated Codelivery of AntimiR-10b and AntimiR-21 for Achieving Triple Negative Breast Cancer Therapy. ACS Nano 9, 2290–2302 (2015). 10.1021/nn507465d

20. C. Chen, S. Zhao, A. Karnad, and J. W. Freeman, The biology and role of CD44 in cancer progression: therapeutic implications. J. Hematol. Oncol. 11, 64 (2018). 10.1186/s13045-018-0605-5

21. Y. Huang, J. Yuan, H. Ding, Y. Song, and G. Qian, Design, synthesis and antitumor activity of a novel PEG-A6-conjugated irinotecan derivative. Bioorg. Med. Chem. Lett. 126847 (2019). 10.1016/j.bmcl.2019.126847

22. S. A. Ghamande, M. H. Silverman, W. Huh, K. Behbakht, G. Ball, et al., A phase 2, randomized, double-blind, placebo-controlled trial of clinical activity and safety of subcutaneous Å6 in women with asymptomatic CA125 progression after first-line chemotherapy of epithelial ovarian cancer. Gynecol. Oncol. 111, 89–94 (2008). 10.1016/j.ygyno.2008.06.028

23. M. Finlayson, Modulation of CD44 activity by A6-peptide. 6, 1–8 (2015). 10.3389/fimmu.2015.00135

24. R. Devulapally, N. M. Sekar, T. V. Sekar, K. Foygel, T. F. Massoud, et al., Polymer nanoparticles mediated codelivery of AntimiR-10b and AntimiR-21 for achieving triple negative breast cancer therapy. ACS Nano 9, 2290–2302 (2015). 10.1021/nn507465d

25. J. Mosafer and M. Teymouri, Comparative study of superparamagnetic iron oxide/doxorubicin co-loaded poly (lactic-co-glycolic acid) nanospheres prepared by different emulsion solvent evaporation methods. Artif. Cells, Nanomedicine Biotechnol. 46, 1146–1155 (2018). 10.1080/21691401.2017.1362415

26. R. Gupta, Firefly Luciferase Mutants as Sensors of Proteome Stress Rajat Gupta. (2012).

27. R. Warin, D. Xiao, J. A. Arlotti, A. Bommareddy, and S. V Singh, Inhibition of human breast cancer xenograft growth by cruciferous vegetable constituent benzyl isothiocyanate. Mol. Carcinog. 49, 500–507 (2010). 10.1002/mc.20600

28. J. P. Miller, C. Egbulefu, J. L. Prior, M. Zhou, and S. Achilefu, Gradient-Based Algorithm for Determining Tumor Volumes in Small Animals Using Planar Fluorescence Imaging Platform. Tomography 2, 17–25 (2016). 10.18383/j.tom.2016.00100

29. J. Sun, Y. Liu, Y. Chen, W. Zhao, Q. Zhai, et al., Doxorubicin delivered by a redox-responsive dasatinib-containing polymeric prodrug carrier for combination therapy. J. Control. Release 258, 43–55 (2017). 10.1016/j.jconrel.2017.05.006

30. Y. Ji, X. Yang, Z. Ji, L. Zhu, N. Ma, et al., DFT-Calculated IR Spectrum Amide I, II, and III Band Contributions of N-Methylacetamide Fine Components. ACS Omega 5, 8572–8578 (2020). 10.1021/acsomega.9b04421

31. Y. Nakamura, A. Mochida, P. L. Choyke, and H. Kobayashi, Nanodrug Delivery: Is the Enhanced Permeability and Retention Effect Sufficient for Curing Cancer? Bioconjug. Chem. 27, 2225–2238 (2016). 10.1021/acs.bioconjchem.6b00437

32. S. A. A. Rizvi and A. M. Saleh, Applications of nanoparticle systems in drug delivery technology. Saudi Pharm. J. 26, 64–70 (2018). 10.1016/j.jsps.2017.10.012

33. B. S. Zolnik, S. T. Stern, J. M. Kaiser, Y. Heakal, J. D. Clogston, et al., Rapid distribution of liposomal short-chain ceramide in vitro and in vivo. Drug Metab. Dispos. 36, 1709–1715 (2008). 10.1124/dmd.107.019679

34. J. D. Clogston and A. K. Patri, Zeta Potential Measurement. in Charact. Nanoparticles Intend. Drug Deliv., edited by S. E. McNeil (Humana Press, Totowa, NJ, 2011), pp. 63–70

35. L. J. Mohan, L. McDonald, J. S. Daly, and Z. Ramtoola, Optimising PLGA-PEG Nanoparticle Size and Distribution for Enhanced Drug Targeting to the Inflamed Intestinal Barrier. Pharmaceutics 12, (2020). 10.3390/pharmaceutics12111114

36. S. Honary and F. Zahir, Effect of zeta potential on the properties of nano-drug delivery systems - A review (Part 1). Trop. J. Pharm. Res. 12, 255–264 (2013). 10.4314/tjpr.v12i2.19

37. P. Kattel, S. Sulthana, J. Trousil, D. Shrestha, D. Pearson, et al., Effect of Nanoparticle Weight on the Cellular Uptake and Drug Delivery Potential of PLGA Nanoparticles. ACS Omega 8, 27146–27155 (2023). 10.1021/acsomega.3c02273

38. R. K. Mittapalli, C. Guo, S. K. Drescher, and D. Yin, Oncology dose optimization paradigms: knowledge gained and extrapolated from approved oncology therapeutics. Cancer Chemother. Pharmacol. 90, 207–216 (2022). 10.1007/s00280-022-04444-0

39. D. C. Arruda, T. D. de Oliveira, P. H. F. Cursino, V. S. C. Maia, R. Berzaghi, et al., Inhibition of melanoma metastasis by dual-peptide PLGA NPS. Biopolymers 108, (2017). 10.1002/bip.23029

40. R. A. Davey, T. J. Longhurst, M. W. Davey, L. Belov, R. M. Harvie, et al., Drug resistance mechanisms and MRP expression in response to epirubicin treatment in a human leukaemia cell line. Leuk. Res. 19, 275–282 (1995). 10.1016/0145-2126(94)00159-8

41. C.-P. Day, G. Merlino, and T. Van Dyke, Preclinical mouse cancer models: a maze of opportunities and challenges. Cell 163, 39–53 (2015). 10.1016/j.cell.2015.08.068

42. B. Guo, J. Wei, J. Wang, Y. Sun, J. Yuan, et al., CD44-targeting hydrophobic phosphorylated gemcitabine prodrug nanotherapeutics augment lung cancer therapy. Acta Biomater. 145, 200– 209 (2022). 10.1016/j.actbio.2022.04.016

43. M. P. Doogue and T. M. Polasek, The ABCD of clinical pharmacokinetics. Ther. Adv. Drug Saf. 4, 5–7 (2013). 10.1177/2042098612469335

44. S. C. Turfus, R. Delgoda, D. Picking, and B. J. Gurley, Pharmacokinetics. in Pharmacogn. Fundam. Appl. Strateg., edited by S. Badal and R. B. T.-P. Delgoda (Academic Press, Boston, 2017), pp. 495–512

45. S. Acharya and S. K. Sahoo, PLGA nanoparticles containing various anticancer agents and tumour delivery by EPR effect. Adv. Drug Deliv. Rev. 63, 170–183 (2011). http://10.0.3.248/j.addr.2010.10.008

46. Y. Zhong, T. Su, Q. Shi, Y. Feng, Z. Tao, et al., Co-administration of irgd enhances tumor-targeted delivery and anti-tumor effects of paclitaxel-loaded plga nanoparticles for colorectal cancer treatment. Int. J. Nanomedicine 14, 8543–8560 (2019). 10.2147/IJN.S219820

47. S. Rezvantalab, N. I. Drude, M. K. Moraveji, N. Güvener, E. K. Koons, et al., PLGA-based nanoparticles in cancer treatment. Front. Pharmacol. 9, 1–19 (2018). 10.3389/fphar.2018.01260

48. D. E. 3rd Owens and N. A. Peppas, Opsonization, biodistribution, and pharmacokinetics of polymeric nanoparticles. Int. J. Pharm. 307, 93–102 (2006). 10.1016/j.ijpharm.2005.10.010

49. J. Li, C. Chen, and T. Xia, Understanding Nanomaterial-Liver Interactions to Facilitate the Development of Safer Nanoapplications. Adv. Mater. 34, e2106456 (2022). 10.1002/adma.202106456

50. J. Huang, L. Bu, J. Xie, K. Chen, Z. Cheng, et al., Effects of Nanoparticle Size on Cellular Uptake and Liver MRI with Polyvinylpyrrolidone-Coated Iron Oxide Nanoparticles. ACS Nano 4, 7151–7160 (2010). 10.1021/nn101643u

51. S. Milewska, A. Sadowska, N. Stefaniuk, I. Misztalewska-Turkowicz, A. Z. Wilczewska, et al., Tumor-Homing Peptides as Crucial Component of Magnetic-Based Delivery Systems: Recent Developments and Pharmacoeconomical Perspective. Int. J. Mol. Sci. 25, (2024). 10.3390/ijms25116219

52. M. Agarwal, A. K. Sahoo, and B. Bose, Receptor-Mediated Enhanced Cellular Delivery of Nanoparticles Using Recombinant Receptor-Binding Domain of Diphtheria Toxin. Mol. Pharm. 14, 23–30 (2017). 10.1021/acs.molpharmaceut.6b00480

53. S. Tang, P. Wen, K. Li, J. Deng, and B. Yang, Tumor targetable and pH-sensitive polymer nanoparticles for simultaneously improve the Type 2 Diabetes Mellitus and malignant breast cancer. Bioengineered 13, 9754–9765 (2022). 10.1080/21655979.2022.2060721

54. R. Yazdian-Robati, E. Amiri, H. Kamali, A. Khosravi, S. M. Taghdisi, et al., CD44-specific short peptide A6 boosts cellular uptake and anticancer efficacy of PEGylated liposomal doxorubicin in vitro and in vivo. Cancer Nanotechnol. 14, 1–13 (2023). 10.1186/s12645-023-00236-0

55. R. S. Piotrowicz, B. B. Damaj, M. Hachicha, F. Incardona, S. B. Howell, et al., A6 peptide activates CD44 adhesive activity, induces FAK and MEK phosphorylation, and inhibits the migration and metastasis of CD44-expressing cells. Mol. Cancer Ther. 10, 2072–2082 (2011). 10.1158/1535-7163.MCT-11-0351

56. Y.-Q. Huang, J.-D. Yuan, H.-F. Ding, Y.-S. Song, G. Qian, et al., Design, synthesis and antitumor activity of a novel PEG-A6-conjugated irinotecan derivative. Bioorg. Med. Chem. Lett. 30, 126847 (2020). 10.1016/j.bmcl.2019.126847

57. S. Murugan, P. Vadevoo, S. Gurung, H. Lee, G. R. Gunassekaran, et al., Peptides as multifunctional players in cancer therapy. (2023). 10.1038/s12276-023-01016-x

58. J. Regberg, A. Srimanee, and Ü. Langel, Applications of cell-penetrating peptides for tumor targeting and future cancer therapies. Pharmaceuticals 5, 991–1007 (2012). 10.3390/ph5090991

59. B. Yameen, W. Il Choi, C. Vilos, A. Swami, J. Shi, et al., Insight into nanoparticle cellular uptake and intracellular targeting. J. Control. Release 190, 485–499 (2014). 10.1016/j.jconrel.2014.06.038

60. M. Mahmoudi, J. Meng, X. Xue, X. J. Liang, M. Rahman, et al., Interaction of stable colloidal nanoparticles with cellular membranes. Biotechnol. Adv. 32, 679–692 (2014). 10.1016/j.biotechadv.2013.11.012

61. R. Ferner and J. Aronson, Susceptibility to adverse drug reactions. Br. J. Clin. Pharmacol. 85, 2205–2212 (2019). 10.1111/bcp.14015

62. R. Mattioli, A. Ilari, B. Colotti, L. Mosca, F. Fazi, et al., Doxorubicin and other anthracyclines in cancers: Activity, chemoresistance and its overcoming. Mol. Aspects Med. 93, 101205 (2023). 10.1016/j.mam.2023.101205

63. J. Lee, M. K. Choi, and I. S. Song, Recent Advances in Doxorubicin Formulation to Enhance Pharmacokinetics and Tumor Targeting. Pharmaceuticals 16, 1–32 (2023). 10.3390/ph16060802

64. M. S. Razavi, A. Abdollahi, A. Malek-Khatabi, N. M. Ejarestaghi, A. Atashi, et al., Recent advances in PLGA-based nanofibers as anticancer drug delivery systems. J. Drug Deliv. Sci. Technol. 85, 104587 (2023). 10.1016/j.jddst.2023.104587

65. M. Zhang and H. Xu, Peptide-assembled nanoparticles targeting tumor cells and tumor microenvironment for cancer therapy. Front. Chem. 11, 1–15 (2023). 10.3389/fchem.2023.1115495.

66. Agudelo, D., Bourassa, P., Bérubé, G., & TajmiR-Riahi, H. A. (2014). Intercalation of antitumour drug doxorubicin and its analogue by DNA duplex: structural features and biological implications. International journal of biological macromolecules, 66, 144–150.

67. Chen, L., Zhao, X., Wang, J., & Xu, Y. (2023). Reciprocal regulation of ABCB1 and miR-21 in cancer drug resistance: Implications for targeted therapy. Journal of Cancer Research and Clinical Oncology, 149(6), 1587–1 598.

68. Gong, X., Zhang, H., & Li, L. (2018). Role of miR-21 in multidrug resistance of MDA-MB-231 breast cancer cells via regulation of ABCB1. Molecular Medicine Reports, 18(6), 5734–5740.

69. Zhang, L., Li, X., & Zhao, Y. (2021). miR-21 upregulates ABCB1 expression and contributes to drug resistance in triple-negative breast cancer. Oncology Reports, 46(4), 1919–1 930.

70. Wang, J., Zhang, R., & Wei, Q. Wang, J., Zhang, R., & Wei, Q. (2022). The role of miR-21 in regulating ABCB1 and its effect on drug resistance in MDA-MB-231 cells. Frontiers in Oncology, 12, 765634

